# The Main protease (M^pro^) from SARS-CoV-2 triggers plasma clotting *in vitro* by activating coagulation factors VII and FXII

**DOI:** 10.1101/2024.09.05.611400

**Authors:** Anna Pagotto, Federico Uliana, Giulia Nordio, Andrea Pierangelini, Laura Acquasaliente, Maria Ludovica Macchia, Massimo Bellanda, Barbara Gatto, Giustina De Silvestro, Piero Marson, Paolo Simioni, Paola Picotti, Vincenzo De Filippis

**Affiliations:** Dept. of Pharmaceutical and Pharmacological Sciences, University of Padua, Padua, Italy; Department of Biology, Institute of Molecular Systems Biology, ETH Zürich, Zürich, Switzerland; Department of Biology, Institute of Biochemistry, ETH Zürich, Zürich, Switzerland; Dept. of Chemical Sciences, University of Padua, Padua, Italy; Department of Transfusion Medicine, University Hospital of Padova, Padua, Italy; Thrombotic and Haemorrhagic Diseases Unit, Department of Medicine, University of Padua, Padua, Italy

**Keywords:** COVID-19, SARS-CoV-2, Main Protease, thrombosis, proteolysis, proteomics

## Abstract

Although the connection between COVID-19 and coagulopathy has been clear since the early days of SARS-CoV-2 pandemic, the underlying molecular mechanisms remain unclear. Available data support that the burst of cytokines and bradykinin, observed in some COVID-19 patients, sustains systemic inflammation and the hypercoagulant state, thus increasing thrombotic risk. Here we show that the SARS-CoV-2 main protease (Mpro) can play a direct role in the activation of the coagulation cascade. Adding Mpro to human plasma from healthy donors increased clotting probability by 2.5-fold. The results of enzymatic assays and degradomics analysis indicate that Mpro triggers plasma clotting by proteolytically activating coagulation factors zymogens VII and XII at their physiological activation sites, i.e. Arg152-Ile153 bond for FVII and Arg353-Val354 bond for FXII, where FVII and FXII are strategically positioned at the very beginning of the extrinsic or intrinsic pathways of blood coagulation. These findings are not compatible with the substrate specificity of the protease known so far, displaying a prevalence for a Gln-residue in P1 and a hydrophobic amino acid in P2 position. This apparent discrepancy was resolved by High Throughput Protease Screen assay, unveiling an extended, time-dependent, secondary specificity of Mpro for Arg-X bonds, which was further confirmed by Hydrogen-Deuterium Exchange Mass spectrometry analysis of Arg-containing inhibitors binding to Mpro and by enzymatic assays showing that the protease can cleave peptide substrates containing Arg in P1. Overall, integrating biochemical, proteomics and structural biology experiments, we unveil a novel mechanism linking SARS-CoV-2 infection to thrombotic complications in COVID-19.

## 1. INTRODUCTION

Clinical evidence accumulated during the last decades provided new insights for a positive correlation existing between infectious diseases and dramatic thrombotic complications. Indeed, systemic or localized infections increase the risk of thrombosis by about 2–20 times and have been recognized as independent risk factors for thromboembolic diseases, as well as for cardiovascular (e.g., myocardial infarction) and cerebrovascular events (e.g., stroke)^1^. Among the viruses associated with thrombotic complications in humans are exanthematous viruses such as varicella, variola, measles, and vaccinia; arboviruses like dengue virus; ebola virus; and also influenza virus, hepatitis viruses, human immunodeficiency virus, and cytomegalovirus, as earlier reviewed ^2–8^. While no viral proteases have yet demonstrated direct activation of coagulation, the crosstalk between virus infection and pro-coagulant signalling pathways has been elucidated in the cases of Ebola^9^ and Dengue^10^ viruses. It has been observed that during infection, the complex Tissue Factor (TF)/coagulation factor VII (FVII) is activated, fostering a pro-coagulant state. It is thought that the activation of coagulation serves as a host protective mechanism to restrain virus spread, although an excessive activation of coagulation may result in haemorrhagic events due to depletion of circulating coagulation factors, as seen in Ebola and Dengue haemorrhagic fever^3^.

Similarly, the outbreaking viral pneumonia (COVID-19), caused by the beta coronavirus SARS-CoV-2, has dramatically strengthened the observed relationship existing between thrombotic complications and virus infection^11^. Critically ill patients with COVID-19 exhibit a 15% incidence of venous thromboembolism and 30% occurrence of arterial thrombosis, while autopsies revealed an incidence of deep venous thrombosis in 58% of non-survivors COVID-19 patients. These clinical studies identified a positive correlation between the incidence of thrombogenesis and the high mortality observed in severe COVID-19 cases. Indeed, approximately 50% of non-survivors presented a pro-coagulant state, whereas only 7% of survivors were pro-coagulant^11^. Consequently, clinicians adopted an anti-coagulation prophylaxis (mainly based on heparin treatment) to prevent thrombosis in patients with moderate-severity COVID-19. However, this approach was considered not suitable for patients requiring ICU-level care, due to the potential for increased bleeding^12^. While hypercoagulability stands as a hallmark for SARS-CoV-2 infection and test of coagulations are currently employed to prognostic the syndrome^13^, there is still a large demand for the characterization of the mechanisms leading to thrombotic events in patients with severe COVID-19, as this knowledge may represent new targets for safer therapies ^14–17^. Since severe COVID-19 has been associated with a massive release of proinflammatory cytokines^18,19^, and high bradykinin (BK) levels^11^, the two current hypotheses entail the “cytokines storm” ^20,21^ and the “bradykinin storm”^11^. Indeed, cytokines (TNFα, IL-1, IL-6, IL-8) promote the expression of intravascular TF, thus leading to the activation of the coagulation cascade via the activation of FVII ^22^. The impact of cytokines is further heightened by IL-6, which can stimulate the formation of new platelets exhibiting an increased sensitivity to thrombin activation^23^. In addition to this inflammation-driven mechanism, the bradykinin-mediated signaling pathway contributes to the activation of pro-inflammatory and pro-coagulant cytokines, creating a positive vicious feedback loop^24^.

Blood coagulation is the result of concomitant events occurring within the blood vessels. Those include the sequential activation of coagulation factors via proteolytic cleavage of zymogen at conversed sites^25^, typically Arg↓Ile/Val bonds, which finally results in the generation of active thrombin and the conversion of fibrinogen to fibrin^26^. In a previous work^27^, we have demonstrated that subtilisin (50 nM – 2 μM), a serin-protease secreted by *Bacillus subtilis* (a non-pathogenic commensal in human gut microbiota), can clot human blood by converting prothrombin into active thrombin species via limited proteolysis, thus bypassing the canonical activation of the coagulation cascade. By the same token, in this work we probe whether a similar proteolytic mechanism occurs during the development of SARS-CoV-2 infection.

SARS-CoV-2 is a positive-strand RNA coronavirus with an unusually large genome that spans about 30kb and encodes at least 13 open reading frames^28^. Among these, the replicase polyprotein 1ab (R1AB) assumes the crucial role of generating the non-structural proteins forming the replicase-transcriptase complex, essential for the RNA-synthesizing machinery. During viral maturation, two key proteases encoded in the R1AB gene, i.e. the main protease M^pro^ (also known as 3CL protease nsp5) and the papain like PL^pro^ nsp3 protease, cleave the replicase polyprotein R1AB promoting the assembly of the replicase-transcriptase complex that encodes for the four structural proteins, i.e. the envelope (E), membrane (M), spike (S) and nucleocapsid (N) proteins^28^. Aside from the primary targets in R1AB gene, M^pro^ and PL^pro^ are pleiotropic viral factor generating proteolytic cleavages in the host organism that result in the re-organization of the cellular machinery and, in some case, in the evasion from the cell resistance mechanisms^29–32^. Among the proteases encoded in SARS-CoV-2 genome, M^pro^ emerges as a potent antigen in infected individuals and, indeed, anti-M^pro^ antibodies were detected in the sera of COVID-19 patients^33,34^. Even though at the best of our knowledge the M^pro^ protease has yet to be detected in SARS-CoV-2 patient plasma, the mRNA originating from SARS-CoV-2 R1AB gene and encoding for M^pro^ and PL^pro^ proteases has been detected in extra-respiratory districts like blood and faeces, while anti-M^pro^ antibodies were found in the saliva and plasma of SARS-CoV-2 patients, with a positive correlation existing between antibody levels and disease severity ^33,34^. These findings strongly suggest that M^pro^ and other viral constituents circulate in the blood of COVID-19 patients^35,36^, whereby they may interact with host plasma proteins, including coagulation factors which are significantly represented in human plasma. Several studies have already deeply characterized the interaction network between M^pro^ and host proteins in human embryonic kidney and carcinoma colon cell lines^30,36–38^. Nevertheless, M^pro^ direct binding partners in blood serum and potential substrates remain still unexplored. A suggestion for a possible functional role of M^pro^ in the plasma is provided by a modelling study showing a remarkable three-dimensional fold similarity between M^pro^ and FXa and Thrombin^39^, suggesting that M^pro^ could share with these coagulation factors some molecular recognition properties and substrate specificity.

M^pro^ is a 67.6 kDa homodimeric cysteine protease constituted by three domains (domains I,II, and III), with the catalytic dyad His41-Cys145 located in the cleft between domains I and II^40^. The substrate specificity of M^pro^ is characterized by a canonical specificity, with a Gln-residue at P1 and a hydrophobic residue (Leu/Met) at P2 that mirrors the cleavage sites required to release non-structural proteins from R1AB gene expression products ^41–43^. This specificity has been further supported by studies utilizing a combinatorial library of synthetic fluorogenic substrates^44^, as well as by mass spectrometry degradomics studies^30,31,45^, and set the basis for the development of novel M^pro^ inhibitors for treating SARS-CoV-2 infection^46,47^. However, it is noteworthy that the same degradomics studies have revealed that M^pro^ cleavage specificity is not as strict as the canonical definition ^41–43^ and, indeed, M^pro^ can also cleave substrates containing a His-or a Met-residue at P1 position ^30,45^.

In this study, we found that the addition of M^pro^ to human plasma samples significantly enhances the probability of plasma clotting. Using a wide array of biochemical, biophysical and proteomics techniques, we demonstrate that M^pro^ directly cleaves and activates factors VII and XII at their canonical activation sites (i.e., Arg152-Ile153 bond in FVII and Arg353-Val354 bond in FXII), where FVII and FXII are key zymogens strategically positioned at the beginning of the extrinsic or intrinsic pathway of the coagulation cascade. Furthermore, using advanced mass-spectrometry methods, we discovered that (alongside the conventional preference for a Gln at P1 position) the protease can cleave (albeit with lower specificity) peptidyl substrates with an Arg-residue at P1 position. This secondary specificity, combined with the observed pro-coagulant effect of M^pro^ and its potential to circulate in COVID-19 patient bloodstream (see above), supports the hypothesis that M^pro^ can play a direct role in the activation of blood coagulation and the onset of thrombotic complications in SARS-CoV-2 infections.

## 2. METHODS

### 2.1. Reagents

Human coagulation factors Prothrombin, Thrombin, FX, FXa, FVII, FVIIa, FIX, FIXa, FXI, FXIa, FXII and recombinant TF (TF), were purchased from Haematologic Technologies Inc. (Essex Junction, VT, USA); fibrinogen from American Diagnostica (Stamford, CT, USA). Commercial M^pro^ (rcM^pro^) was purchased from Sigma-Aldrich (St. Louis, MO, USA). Chromogenic substrates S2238, S2765, S2366, S2302 were purchased from Chromogenix (Milan, Italy), while MeSO_2_-D-CHA-But-Arg-pNA was from Sigma (Darmstadt, Germany). M^pro^ FRET-based substrate 5-FAM-AVLQSGFRK(DABCYL)K (ProteoGenix, Miami, FL, USA)-Commercial human plasma (STA-System Control N+P, Normal plasma) was purchased from STAGO (Milan, Italy). Egg phosphatidylcholine (PC) and brain phosphatidylserine (PS) were purchased from Avanti Polar Lipids, Inc. (Alabaster, AL, USA). Salts, solvents, and other reagents were of analytical grade and purchased from Merck LifeScience (Darmstadt, Germany).

### 2.2. Protein preparation

In-house produced recombinant M^pro^ (rM^pro^) was purified and characterized as previously described^48^. Recombinant commercial M^pro^ (rcM^pro^) from SIGMA was resolubilised in water containing 1mM DTT, to reach a final concentration of 1mg/ml. Commercial human fibrinogen (America Diagnostica Inc) was desalted on a 5-ml HiTrap desalting column (GE Healthcare, IL, USA), eluted at a constant flow rate (1 ml/min) with 10 mM HEPES pH 7.4, 150 mM NaCl (HBS). The material eluted in correspondence of the major chromatographic peak was collected and used for subsequent analyses. Fibrinogen concentration was determined spectrophotometrically at 280 nm, using a molar absorptivity of 513.400 M^-1^·cm^-1^.

### 2.3. Human plasma preparation

Platelet poor plasma (PPP) was prepared from freshly withdrawn, citrated blood samples taken from 21 healthy donors (twelve men and nine women, 21–60 years of age). Donors gave their written informed consent to participate in this study, which was approved by the Institutional Ethics Committee of the Padua University Hospital and all methods were performed in accordance with the relevant guidelines and regulations. PPP was prepared by centrifugation of whole blood at 1,500 g for 15 min, at room temperature, without brake, as recommended by the Clinical Laboratory Standards Institute guidelines^49^. For commercial plasma samples preparation, citrated lyophilized normal human plasma was reconstituted with distilled water following the manufacturer’s instructions.

### 2.4. Liposomes preparation

Liposomes containing phosphatidyl choline (PC) and phosphatidyl serine (PS) were obtained by the extrusion method, as detailed elsewhere^50^. The final liposomes solution (12 μM) was composed by PC and PS (50:50 molar ratio) unilamellar vesicles of 100±30 nm diameter, as determined by Dynamic Light Scattering (DLS), using a Zetasizer Nano-S instrument (Malvern Instruments, UK).

### 2.5. Turbidimetric assays

Fibrin generation by rcM^pro^ was probed on re-calcified citrated plasma diluted 1:2 with HBS or on a solution of freshly desalted fibrinogen (0.15 mg/ml). To assess the correct CaCl_2_ amount to be added to citrate plasma, a re-calcification scouting was run, and the maximum CaCl_2_ concentration which did not trigger plasma clotting within 1 hour was selected. Plasma samples or fibrinogen were incubated with increasing concentration of rcM^pro^, at 37±0.1°C, or the correspondent higher amount of added DTT as a control, and the clot formation was followed by continuously recording the apparent absorbance of the solution at 350 nm, using a V-630 Jasco (Tokyo, Japan) spectrophotometer or a Victor Nivo Multiplate Reader (PerkinElmer, Waltham, MA, USA). Positive controls were performed with human α-thrombin (2nM). The resulting clotting curves were analysed to extract the values of *t_c_,* and Δ*A_max_*, where *t_c_* is the clotting time and it is calculated from the intercept point of the tangent to the maximal slope of the curve with the time axis; *ΔA_max_*is the maximal absorbance change when fibrin generation is complete^27^. Statistical difference in the probability of plasma clotting after rcM^pro^ treatment, was assessed by performing turbidimetric assay on plasma samples from 20 healthy donors. Each clotting curve was evaluated for the presence of a sigmoidal curve indicating a clotting event, and the clotting time was accordingly calculated. The probability of clotting within 1h of assay was estimated using the Kaplan Meier method and the survival curves were compared by log rank test^51^ using the R-packages “survival” and “survminer” ^52,53^.

### 2.6. Electrophoretic analysis of plasma clot

After 1-hour incubation at 37°C of 1:2 diluted plasma with rM^pro^ (200 nM), the plasma clot was collected, washed with HBS buffer and resuspended with GdnCl 8M (incubated for 1h at 37°C). Albumin was then removed by a treatment with EtOH to a final concentration of 42% for 1h at 4°C and following centrifugating at 16,000g for 45min at 4°C. After removing the supernatant, the pellet was then resuspended with SDS sample loading buffer and analysed by reducing SDS-PAGE (Bolt Bis-TRIS 4-12% precast gel) and Coomassie staining (Simply Blue SafeStain, Invitrogen). For comparison, commercial fibrinogen (Fbg) was also loaded. The typical α-, β– and y-chain of Fbg are indicated by arrows, along with the y-chain dimer.

### 2.7. Coagulation factors activation assays

Activation of coagulation factors zymogens by M^pro^ was assessed by incubating at 37±0.1°C the zymogen in the absence (negative control) and presence of rcM^pro^ and monitoring at the same temperature the release of *p*-nitroaniline (pNA) from the corresponding specific chromogenic substrate by measuring the absorbance increase at 405 nm (□^M^_405nm_ = 9,920 M^-1^·cm^-1^). For each zymogen, the experimental details are reported in the following. Prothrombin (10nM) and FX (10nM) were incubated with rcM^pro^ at a E:S ratio of 10:1 (mol/mol) in HBS buffer, containing 0.1% (w/v) PEG_8000_ (HBS-PEG) and 5mM CaCl_2_. After 3-h incubation, the reaction mixture was added with α-thrombin chromogenic substrate S2238 (20 μM) or with FXa chromogenic substrate S2765 (100 μM). Activation of prothrombin with 1nM FXa in the presence of 100μM PCPS (50:50) and activation of FX with 0.2nM FVIIa in the presence of 2nM TF were performed as positive controls. FVII (200nM) was incubated for increasing periods of time in HBS-PEG with 5mM CaCl_2_, at 37±0.1°C, alone or with rcM^pro^ at an E:S ratio of 10:1 (mol/mol). At time points (i.e., 0, 45, 90, 180 min), aliquots of FVII from the reaction mixture (10 nM final concentration) were added to FVIIa-specific chromogenic substrate MeSO_2_-D-CHA-But-Arg-pNA (300 μM) in the presence of TF (100 nM). To calculate the concentration of the FVIIa generated by rcM^pro^, at each time point, the absorbance signal obtained from FVII in the presence of TF alone (reference signal) was subtracted from the absorbance signal obtained in the presence of rcM^pro^, and the initial rate (v_0_) of pNA release was calculated. The concentration of FVIIa was then estimated from a FVIIa titration curve in the presence of 100nM TF. FIX (100nM) was incubated with rcM^pro^ at an E:S ratio of 1:1 (mol/mol) in HBS-PEG with 3mM CaCl_2_. FXI activation was determined indirectly by activation of FX. Hence, after 3-h incubation, the reaction mixture was added with FX (10nM) and FXa chromogenic substrate S2765 (100 μM). Activation of FX with 40nM FIXa in the presence of 20μM PCPS (50:50) was performed as a positive control. FXI (1nM) was incubated with rcM^pro^ at a E:S ratio of 1:1 (mol/mol) in HBS-PEG with 3mM CaCl_2_. After 3-h incubation, the reaction mixture was added with FXIa chromogenic substrate S2366 (100 μM). Cleavage of S2366 by FXIa (0.1nM) was performed as a positive control. FXII (100nM) was treated with rcM^pro^ at an E:S ratio of 1:1 (mol/mol) in HBS with 5mM CaCl_2_ and added with FXIIa chromogenic substrate S2302 (300 μM) without previous incubation. Activation of FXII in the presence of 40μM PCPS (50:50) was performed as a positive control. Activation of FXII by rcM^pro^ was compared to BSA at the same E:S ratio.

The effect of rcM^pro^ on the anticoagulant factor Antithrombin III (ATIII) was tested by incubating for 1h at 37°C ATIII with rcM^pro^ in a 1:1 molar ratio (40nM). This solution was further incubated with α-thrombin (1nM) for 3h at 37°C in HBS-PEG, 3mM CaCl_2_. α-Thrombin residual activity, after ATIII incubation, in the absence and presence of rcM^pro^, was tested using the S2238 chromogenic substrate.

### 2.8. HTPS analysis

In the High-throughput native microscale protease screen (HTPS), a standardized native cell lysate is proteolyzed with the studied protease. The protease-generated peptides are collected, analyzed by MS, and the identified substrate peptides are analyzed to retrieve activity, specificity and cleavage entropy data. The detailed workflow is described in *Uliana et al.*^54^.

*Sample preparation*. For HTPS characterization of rcM^pro^, a variation of the protocol for cysteine protease has been developed^55^: native cell lysate in 20mM Ammonium bicarbonate pH 7.8 (50µg, 0.5mg/ml), 5mM L-cysteine, 1mM EDTA was incubated with rcM^pro^ (5 µg, E:S 1:10) for 5min, 30min, 4h and 24h at 37±0.1°C under agitation (1,400g) in a filtered 1.5ml tube (10-kDa cutoff) from Sartorius (Göttingen, Germany), according to the filter-aided sample preparation (FASP) protocol for proteomic analysis^56^. Peptides were collected by 15-min centrifugation 10,000g followed by one wash with 100µl of MS-grade water to increase peptide recovery. After cleanup using C18 StageTips, dried peptides were dissolved in 2% acetonitrile-0.1% formic acid before the analysis by MS in the data-dependent acquisition (DDA) mode. Samples were prepared in triplicate.

*MS acquisition.* LC-MS/MS was performed on an Orbitrap QExactive+ mass spectrometer (Thermo-Fisher, Waltham, MA, USA) coupled to an EASY-nLC-1000 liquid chromatography system (Thermo-Fisher). Peptides were separated using a reverse phase column (75□µmx400□mm New Objective, in-house packed with ReproSil Gold 120 C18, 1.9 µm, Dr. Maisch GmbH) across 180□min gradient from 3% to 25% B in 160 min and from 25% to 40% B in 20 min (buffer A: 0.1% (v/v) formic acid; buffer B: 0.1% (v/v) formic acid, 95% (v/v) acetonitrile). The DDA acquisition mode was set to perform one MS1 scan followed by a maximum of 20 scans (TOP20) with MS1 scans (R=70,000 at 400□m/z, AGC=3e6 and maximum IT=64ms), HCD fragmentation (NCE=25%), isolation windows (1.4m/z) and MS2 scans (R=35’000 at 400□m/z, AGC=2e5 and maximum IT=55ms). A dynamic exclusion of 30s was applied and charge states lower than two and higher than seven were rejected for the isolation.

*Data analysis.* DDA data were searched with MaxQuant software package (version 1.5.2.8) using HTPS_DB.fasta database (2’557 entries). The search was performed with digestion mode set to unspecific, maximal peptide length to 40 AA and only acetylation N termini and methionine oxidation as variable modifications. First search peptide mass tolerance was 20 ppm and main search peptide mass tolerance was 4/5ppm. MS/MS match tolerance was set to 20ppm. Target decoy approach was used to control FDR which was set to 0.01 at peptide and PSM level. Protease substrate specificity analysis was performed in R (version 3.4.3) using the workflow deposited on Github (https://github.com/anfoss/HTPS_workflow, https://doi.org/10.5281/zenodo.4484341) under MIT license.

### 2.8. Chemical synthesis

The peptide substrates (FTRLQSLEN and FTRLRSLEN) were synthesized by the manual solid-phase method using the 9 fluorenylmethyloxycarbonyl (Fmoc)/t butyl (tBu) strategy^57^. After peptide chain assembly, resin cleavage and removal of side-chian protecting groups, the crude peptides were fractionated by RP-HPLC on a (4.6 ×□250□mm, 5μm granulometry, 300□Å pores size) C18 analytical column (Grace-Vydac, Hesperia, CA, USA), equilibrated with 0.1% (v/v) aqueous TFA and eluted with a linear acetonitrile-0.078% (v/v) TFA gradient. The peptide material eluted in correspondence of the major chromatographic peaks was collected, lyophilized, and analyzed by HR-MS, yielding mass values in agreement with the theoretical mass within 2ppm accuracy.

Peptide samples (10 µM) in HBS were incubated at 37°C in the presence of rcM^pro^, at a protease:peptide molar ratio of 1:40. At given time points (0 min, 1 hour, 2 hours, 4 hours, 7 hours, 16 hours) aliquots (5 pmol) were taken, acid quenched with 0.1% (v/v) aqueous TFA, and then analysed by LC-MS. Quantitative determination of intact peptides and proteolytic fragments (FTRLQSLEN, FTRLQ, SLEN, FTRLRSLEN, FTRLR, SLEN) was performed by integrating the area under the chromatographic peaks, using the Jasco ChromNav 2.0 software.

### 2.9. Inhibition of M^pro^ proteolytic activity by thrombin inhibitors Argatroban and PPACK

The inhibition of rcM^pro^ by thrombin and FXa inhibitors Argatroban or PPACK was measured through an *in vitro* fluorogenic assay, using the fluorogenic rcM^pro^ substrate 5-FAM-AVLQSGFRK(DABCYL)K. rcM^pro^ (50nM) was incubated with Argatroban or PPACK (7μM) in HBS-PEG for 2h, at 37°C. After the addition of 5-FAM-AVLQSGFRK(DABCYL)K (1.25μM), the emission of the released peptidyl 5-FAM fluorophore was measured at 530nm. The residual proteolytic activity of rcM^pro^ was determined as the ratio v_i_/v_0_, where v_i_ and v_0_ are the initial velocities of M^pro^-catalyzed substrate hydrolysis in the presence or in the absence of the inhibitor.

To calculate the inhibition constant (K_i_) of PPACK, M^pro^ residual activity was measured at increasing concentrations of PPACK after 10-min incubation at 37°C. The ratio v_i_/v_0_ was plotted as a function of PPACK concentration and data points were fitted with equation 1, describing the slow-tioght binding inhibition model ^58^ to yield the apparent inhibition constant (K_i_ ^app^):

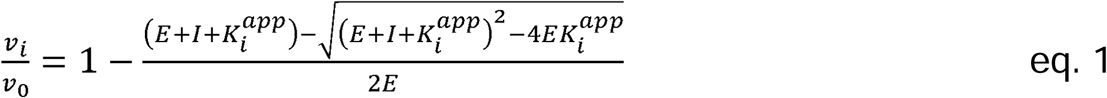

where E is the rcM^pro^ concentration, and I is the PPACK concentration. The equilibrium inhibition constant (K_i_) was then derived from K_i_ ^app^ using equation 2, according to the competitive inhibition model:

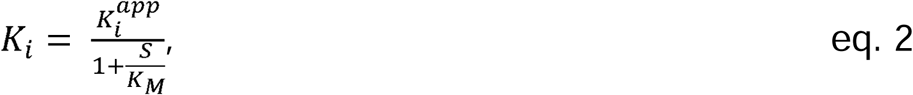

where S is the substrate concentration, and K_M_ is the Michaelis-Menten constant of rcM^pro^ for 5-FAM-AVLQSGFRK(DABCYL)K, calculated as 20 μM.

### 2.10. Computational methods

*Molecular docking*. Molecular docking was performed with HPEPDOCK web server ^52^, starting from the inhibitor-free M^pro^ structure, after removal water molecules (6y2e.pdb) ^35^ and the (D)-Phe-Pro-Arg peptide, as the peptidyl moiety of the irreversible inhibitor PPACK. The software generated 3D structure models for the given sequences of peptide using the implemented MOPEP software, which considers peptide flexibility. Simulations were run with default parameters. One hundred poses were generated and ranked according to the CAPRI criteria ^53^. The pose with the most favourable energy score was selected and used for data analysis.

*Electrostatics*. Electrostatic potential calculations were performed using the APBS software run on the crystallographic structure of M^pro^ (6y2e.pdb) ^35^. A solvent dielectric constant of 78.14 and a protein dielectric constant of 2.0 at 310 K in 150 mM NaCl were used. Final electrostatic maps were constructed by subtracting the protein self-energies from the calculated map using the DXMATH utility in APBS. Images were generated with PyMOL vs. 1.3 (DeLano Scientific, San Diego, CA, USA)

### 2.11. HDX-MS Local analysis

*Sample preparation and LC-MS.* HDX-MS experiment was performed following the procedure described in Peterle et *al*.^59^ and Acqusaliente et *al.*^57^. Briefly, samples were acquired in triplicate using a Waters Xevo G2S mass spectrometer (Waters, Milford, MA, USA) equipped an ACQUITY UPLC M-class chromatographic system (Waters). All sample handling was carried out using a Leap HDX PAL autosampler (Leap Technologies, Carrboro, NC, USA). rcM^pro^ (25μM) was incubated alone or with PPACK (2mM) in PBS for 30min at 20°C. At each incubation time, deuterium labeling was initiated by a 20-fold dilution of the sample with D_2_O (95% D_2_O in 20 mM sodium phosphate pH 7.16, 150 mM NaCl). At given time points (i.e., 15 sec, 1 min, 5 min, 30 min, 2 hours) the labeling reaction was quenched at 1°C by addition of an equal volume of quenching buffer (200 mM sodium phosphate, 1.5 M guanidine hydrochloride (Gdn-HCl), 1 mM Tris(2-carboxyethyl)phosphine hydrochloride (TCEP), adjusted to pH 1.99) to reach a final pH of 2.46. The pH and pD values were measured at 25 °C, using a Mettler-Toledo (Columbus, OH, USA) mod. FiveEasy Plus pH-meter. Quenched samples were immediately injected into the LC system for online pepsin digestion using an in-house prepared immobilized pepsin column, thermostated at 15°C and equilibrated with an isocratic flow of 0.23% (v/v) formic acid in water at a flow rate of 100μl/min. The resulting peptic fragments were online trapped on an Acquity UPLC BEH C18 VanGuard Pre-column (130Å, 1.7µm, 2.1mmx5mm) and separated using an Acquity UPLC BEH C18 column (130Å, 1.7 µm, 1 mm X 100 mm) with a linear acetonitrile-0.1% (v/v) formic acid gradient from 5% to 60% in 10 min, at a flow rate of 60μl/min. Mass spectra were acquired in the resolution mode (m/z range 50–2000). Unlabeled proteins were prepared as reference samples in the same manner as those that were labeled with deuterium. Each sample was prepared in triplicate.

*Data analysis.* Peptides that were generated from online pepsin digestion were identified from the unlabeled protein samples using the Waters Protein Lynx Global Server 3.0. Only those fragments matching the following criteria were considered: i) a length between 4 and 33 amino acids, ii) at least 2 ion products identified, iii) minimum products per amino acid of 0.2, iv) maximum MH^+^ error tolerance of 6 ppm, and v) the presence of the peptide in at least two of the three peptide identification runs. Those peptides meeting the filtering criteria were further processed by DynamX 3.0 (Waters) to calculate the relative amount of deuteration. Deuterium uptake was obtained by subtracting the centroid mass of the undeuterated form of each peptide from the deuterated form, at each time point, for each condition. Because relative deuterium uptake of individual peptides was compared, no back-exchange correction was performed. The threshold for calling differences in relative deuterium incorporation measurements was set at minimum 0.5 Da.

### 2.12. Identification of M^pro^ cleavages on coagulation factors

The identification of cleavages on coagulation factors was performed using the Terminal Amine Isotopic Labeling of Substrates (TAILS) protocol, which labels the neo-N-termini generated by the protease activity^60^.

*Sample preparation.* Purified commercial human coagulation factors FVII or FXII (4μg) were incubated with rcM^pro^ (1:10 E:S molar ratio) or with a mock treatment in HBS for 3h at 37±0.1°C. After proteolysis, the reactions were terminated by heat-inactivation of the protease at 99°C for 5min. Once cooled down, the samples were denatured with Gdn-HCl (4M final concentration) for 15min at 37°C, reduced with TCEP for 30min at 37°C and alkylated with iodoacetamide (IAA) for 30min at room temperature in the dark. Next, we proceeded with reductive di-methylation of free N-termini by adapting the TAILS protocol^60^. Briefly, samples were incubated with 20 mM formaldehyde and 10 mM sodium cyanoborohydride (NaCNBH_3_) for 16 h at 37°C. The reaction was quenched by 1:5 dilution in 50 mM ammonium bicarbonate pH 7.8 for 4h at r.t., and the labeled protein fragments were digested with trypsin (1:50 ratio (w/w)) for 16 h at 37°C. Tryptic digestion was quenched by acidification with 0.5% formic acid and the samples were desalted following the StageTips C18 protocol^61^. (washing step, 5% acetonitrile containing 0.1% formic acid; elution step, 50% acetonitrile containing 0.1% formic acid). Dried peptides were resuspended in 0.1% formic acid (15μl). Identification and quantification of peptides was performed by liquid chromatography tandem mass spectrometry (LC-MS/MS) operating in Data Dependent Acquisition (DDA) mode or in Parallel Reaction Monitoring (PRM). Samples were prepared in triplicate.

*DDA analysis*. LC-MS/MS was performed on a Orbitrap Eclipse Tribrid mass spectrometer (Thermo-Fisher) coupled to an EASY-nLC2002 (Thermo-Fisher) liquid chromatography system. Peptides were separated using a reverse phase column (75□μmx40□mm New Objective, in-house packed with ReproSil Gold 120 C18, 1.9 μm, Dr. Maisch GmbH) across 60-min linear gradient from 3 to 30% (buffer A: 0.1% [v/v] formic acid; buffer B: 0.1% [v/v] formic acid, 80% [v/v] acetonitrile). The DDA data acquisition mode was set to perform one MS1 scan and multiple MS2 scans in a cycle time of 3s with MS1 identification (R=120,000 at 400□m/z, AGC=200% and maximum IT=100ms, scan range 350-1400), HCD fragmentation (NCE=30%), isolation windows (1.2 m/z) and MS2 identification (R=30,000 at 400□m/z, AGC=200% and maximum IT=54ms, scan range 150-1200). Charge states lower than two and higher than seven were rejected. Acquired spectra were searched using the SpectroMine software (Biognosys, Zurich, Switzerland) against FVII sequence or FXII sequence plus contaminants. The search was performed with the default parameters, changing the digestion specificity to semi-tryptic and adding Lys and N-terminal di-methylation (+29.039125) as a variable modification. The maximum number of modifications per peptide was increased to 3. False discovery rate of <1% was used at peptide level. Only N-terminal di-methylated peptides not identified in any of the mock treatment replicates and identified in at least 2/3 replicates of the rcM^pro^-treated replicates were considered in the analysis.

*PRM analysis.* LC-MS/MS was performed on an Orbitrap Exploris 480 mass spectrometer (Thermo-Fisher) coupled to a Vanquish Neo liquid chromatography system (Thermo-Fisher). Peptides were separated using a reversed phase column (75□μmx400□mm New Objective, in-house packed with ReproSil Gold 120 C18, 1.9μm, Dr. Maisch GmbH) across 60□min linear gradient from 7 to 35% (buffer A: 0.1% [v/v] formic acid; buffer B: 0.1% [v/v] formic acid, 80% [v/v] acetonitrile). MS acquisition of targeted peptide was set up with the combination of one MS1 untargeted scan and MS2 scheduled targeted scan using an isolation window of 2.0 m/z and HCD fragmentation. Manual curated analysis of fragments was performed using Skyline daily^62^. Data analysis of PRM data was performed from the integration of top five most intense fragments and transformed log_2_ values.

## 3. RESULTS

### 3.1. Exogenous rM^pro^ triggers plasma clotting *in vitro*

To assess whether M^pro^ can trigger coagulation in human plasma, we conducted turbidimetric assays by adding the recombinant protease of commercial source (rcM^pro^), or in-house produced (rM^pro^) to plasma samples (**Figure 1A, Figure S1A-B**). Furthermore, we also tested human plasma either of commercial origin or freshly collected from healthy donors, after re-calcification (**Figure S1C**). The resulting clotting curves display three phases: i) a lag phase, corresponding to the time necessary for the elongation of protofibrils from fibrin monomers, ii) a linear rise, resulting from the lateral aggregation of protofibrils, and iii) a plateau, when most of protofibrils have been transformed into fibers^63^. From these curves, we extracted the clotting time (*t_c_*), calculated from the intersect point of the tangent line to the curve at the inflection point with the baseline^64^ and the maximal change in the apparent absorbance at 350nm (Δ*A*).

**Figure 1.**
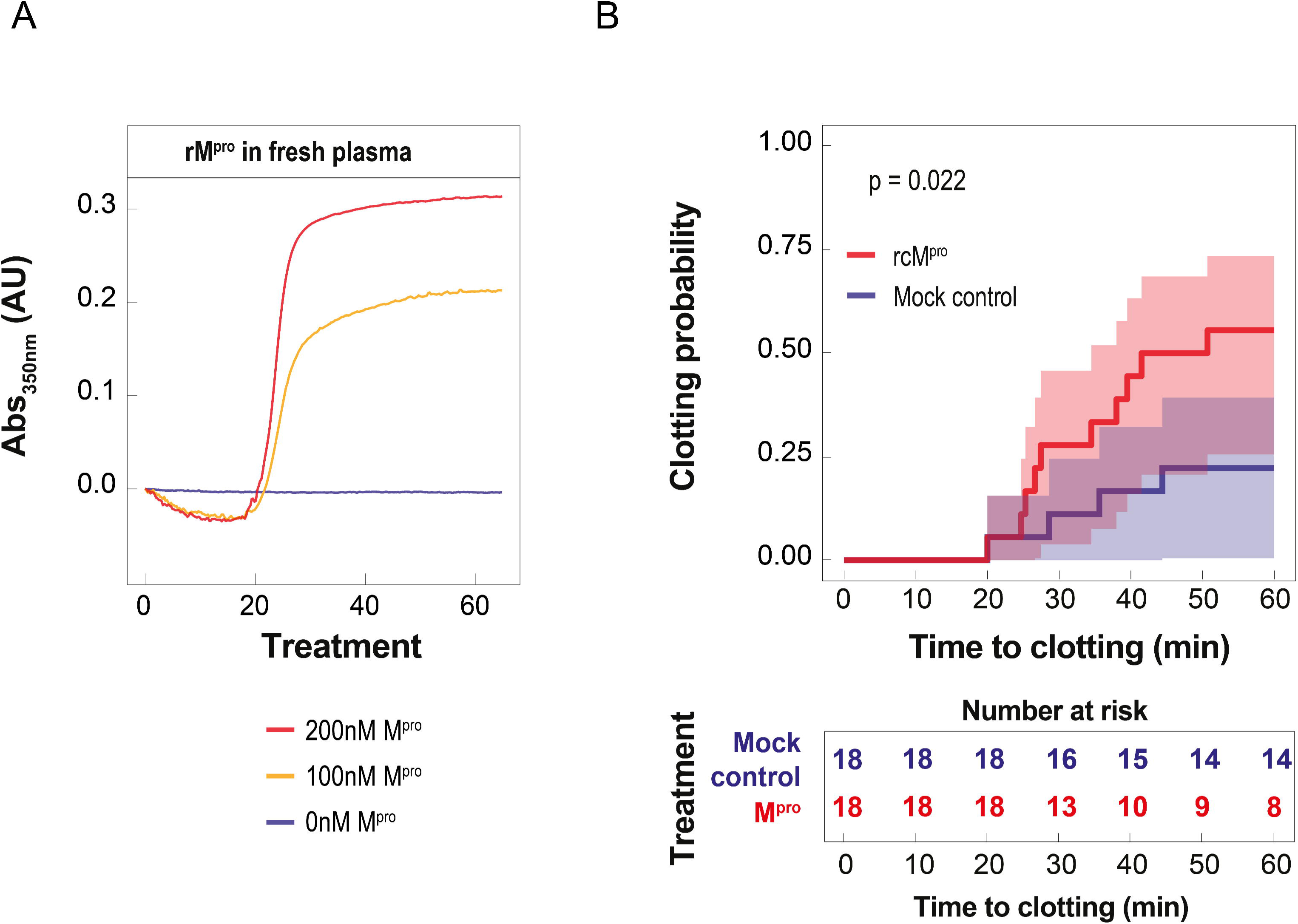
Effect of M^pro^ on fibrin generation in human plasma, monitored by turbidimetric assay. (**A**) Representative clotting curves obtained adding rM^pro^ to human plasma freshly collected from healthy donor. Plasma samples were diluted 1:2 with HBS, re-calcified, and the increase of turbidity at 350 nm was measured over time at 37±0.1°C. The curves resulting from the mean of tree independent experiments are reported in **Figure S1A**. (**B**). Survival curves (Kaplan-Meier) of time to clotting. The clotting probability for plasma incubated with and without M^pro^ (mock control) is compared at different time points. In the lower panel, the number at risk table reports non-clotted plasma. For this analysis the lower M^pro^ concentration triggering coagulation (range from 50 to 500nM) is considered. The statistical analysis is performed using plasma samples from a cohort of 18 healthy donors. For each sample, the clotting curve was manually evaluated for the presence of a sigmoidal trend indicating a clotting event and, accordingly, the clotting time was calculated.

The turbidimetric assay was performed on fresh human plasma samples, collected from healthy donors, after adding 100 or 200 nM rM^pro^. The representative clotting curves, shown in **Figure 1A**, provide clear-cut evidence of clot formation. The presence of fibrin was confirmed by reducing SDS-PAGE (**Figure S1D**), showing the presence of the typical gel bands corresponding to fibrin(ogen) α and β chains, and the y-chain dimer resulting from factor-XIII mediated cross-linking^65^. A t_c_ value of approximately 20 min could be estimated, regardless of rM^pro^ concentration. Furthermore, by doubling the protease concentration, the ΔA_max_ value was increased by about 50%, suggesting that more thrombin is generated at higher rM^pro^ concentration (**Figure 1A**). This conclusion is in keeping with the notion that in human plasma ΔA_max_ is proportional to fibrin clot density^66^ and, ultimately, to thrombin concentration^67^. The same phenotype was observed when repeating the experiment using a commercial recombinant M^pro^ (rcM^pro^) added to freshly prepared plasma (**Figure S1B**), or recombinant in-house produced rM^pro^ added to commercial plasma (**Figure S1C**), indicating that the reagents source does not influence the results.

These initial results prompted us to validate the pro-coagulant effect of M^pro^ in a small cohort of healthy blood donors (n=20). To this aim, we applied non-parametric statistics to estimate the probability of clotting (see the legend to **Figure 1**) in the presence and absence of rcM^pro^ (**Figure S2A,B**). For each clotting curve, the time-dependence of the turbidimetric signal was measured and used to estimate the clotting probability (see the legend to **Figure 1**). Since the identification of clotting time for donors 17 and 20 was difficult to assess (**Figure S2B**), we excluded these two replicates from the statistical analysis. The Kaplan-Meier estimator method^51^ was used to measure at fixed time points (t) the fraction of plasma samples that clotted, with or without rcM^pro^, over one-hour incubation. Despite the inter-individual biological variability, possibly affecting the plasma concentration of some of the coagulation factors tested ^68^, after 60-min incubation with rcM^pro^, a 2.5-fold increase of the clotting probability was detected for the rcM^pro^-treated group (n = 18) compared to the control group (n = 18), with a statistically significant *p*-value of 0.022 (**Figure 1D**, top panel). The greater tendency of plasma to clot in the presence of rcM^pro^ is shown in the table in **Figure 1B** (bottom panel), indicating that at incubation times shorter than 20min all the 18 plasma samples tested remain unclotted, regardless of the presence of rcM^pro^. Conversely, at increasing incubation times, the number of unclotted samples progressively decreases to a greater extent in the rcM^pro^-treated group, compared to the control group. After 1-h incubation, indeed, only eight of the 18 rcM^pro^-treated samples remain unclotted, compared to 14 unclotted samples in the control (rcM^pro^-untreated) group.

Overall, the results of turbidimetric assays provide evidence for a direct procoagulant effect of exogenous M^pro^ on human plasma and encouraged us to further investigate the molecular mechanism(s) by which M^pro^ can induce activation of the coagulation cascade.

### 3.2. M^pro^ selectively activates coagulation factors VII and XII

Fibrin generation is the end-point of the activation of the coagulation cascade in human plasma and the result of a complex and precise balance existing between procoagulant and anticoagulant mechanisms. To clarify the mechanism of M^pro^-induced plasma clotting, we screened *in vitro* the zymogens of pro-coagulant factors (i.e., Fibrinogen, Prothrombin, FVII, FX, FIX, FXI and FXII) for their susceptibility to proteolytic activation by the protease. Moreover, since the observed procoagulant effect of M^pro^ might arise from selective proteolytic degradation/inactivation of anticoagulant factors, we also explored the effect of rcM^pro^ on antithrombin III (ATIII), a major irreversible inhibitor of α-thrombin ^69^.

For the screening of the procoagulant factors activation, the generation of the corresponding active protease was investigated both in the presence and absence of rcM^pro^ by incubating each zymogen with the specific chromogenic substrate releasing *para*-nitroaniline (pNA) (**Figure 2A**). Whereas we did not observe any activation effect by M^pro^ on Prothrombin (ProT or FII), FIX, FX and FXI, nor any inhibition of the effect on ATIII, our results show that M^pro^ treatment significantly (*p-*value < 0.05, Kruskal–Wallis test) increases the release of pNA (log_2_FC >2) for FVII and FXII zymogens (**Figure 2B)** The corresponding progress curves are shown in **Figure 3A,B** for FVII and FXII, and in **Figure S3A-F** for the factors that are not affected by M^pro^.

**Figure 2.**
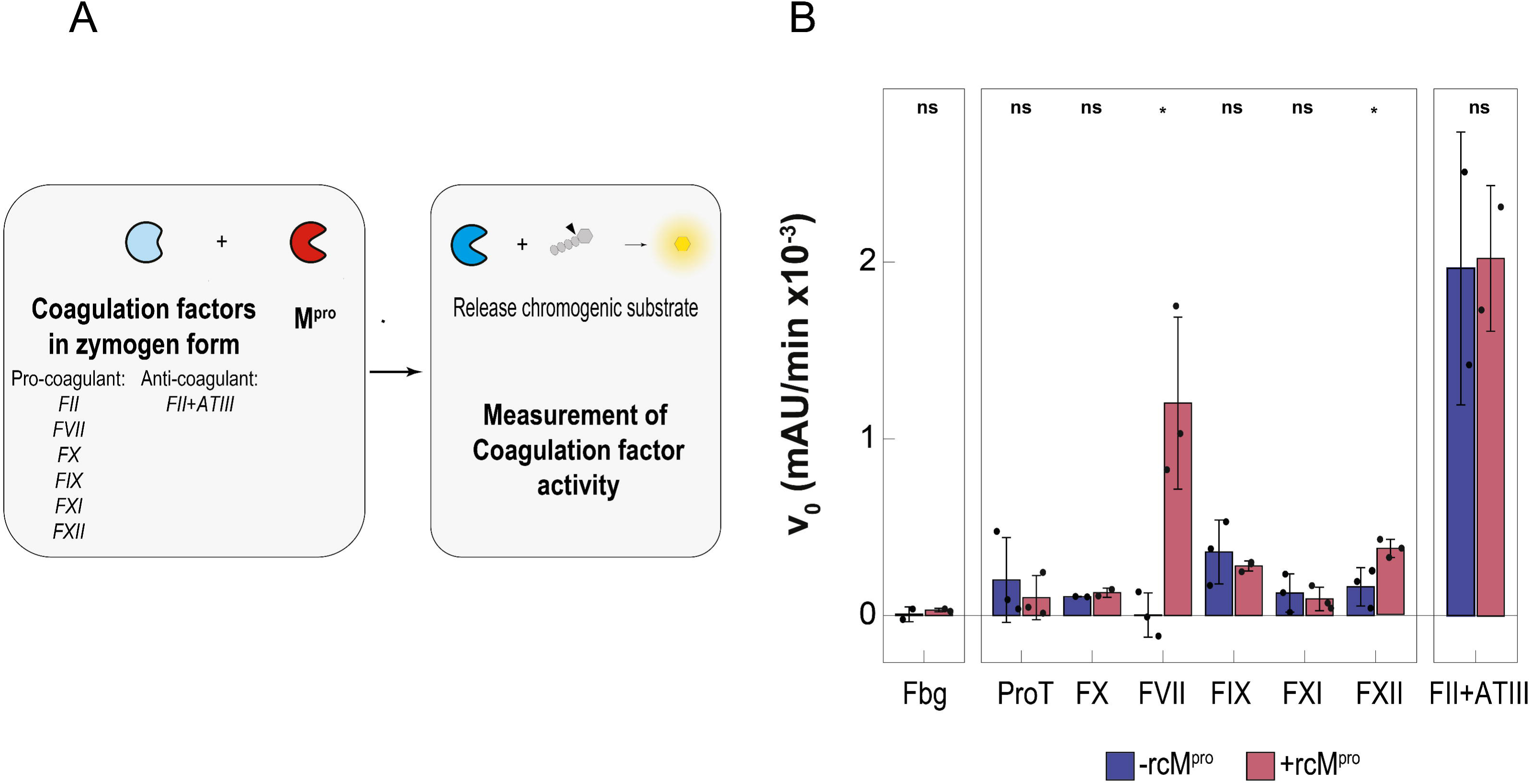
Screening the effect of exogenous M^pro^ on the activation of coagulation factor zymogens. (**A**) **Coagulation factor activation screening workflow**. Coagulation factor zymogens were incubated at 37°C with rcM^Pro^ in HBS-PEG buffer, pH 7.4, for three hours at the specific E:S molar ratio and calcium chloride concentration (see the Methods section, for experimental details). The generation of active proteases was monitored by enzymatic assays, using specific chromogenic substrates and monitoring the release of pNA at 405 nm. FIX activation was indirectly tested by measuring FIXa-mediated activation of FX. The ability of rcM^pro^ to generate fibrin from fibrinogen was assessed by turbidimetric assay (see Methods sections, for experimental details). A positive (i.e., the active coagulation factor added to the chromogenic substrate solution at a concentration identical to that of the zymogen in the proteolysis mixture with rcM^Pro^) and negative (i.e., the coagulation factor zymogen added to the chromogenic substrate solution at a concentration identical to that of the zymogen in the proteolysis mixture with rcM^Pro^) control assay were always run for each test. (**B**) **Screening assays for coagulation factor zymogens activation by M^pro^.** For each zymogen, the bar plot displays the initial rate of pNA release (v_0_) after the treatment with (red bars) or without (blue bars) rcM^Pro^. Data are presented as mean ± standard deviation (SD) and the Kruskal-Wallis test was used for the comparison. Sample size: for ProT/FII, FVII, FIX, FXI, FXII n = 3, for Fbg, FX, FII+ATIII (n = 2). *p ≤ 0.05. The curves are reported in Figure 3A**,B** and **Figure S3A-F**.

**Figure 3.**
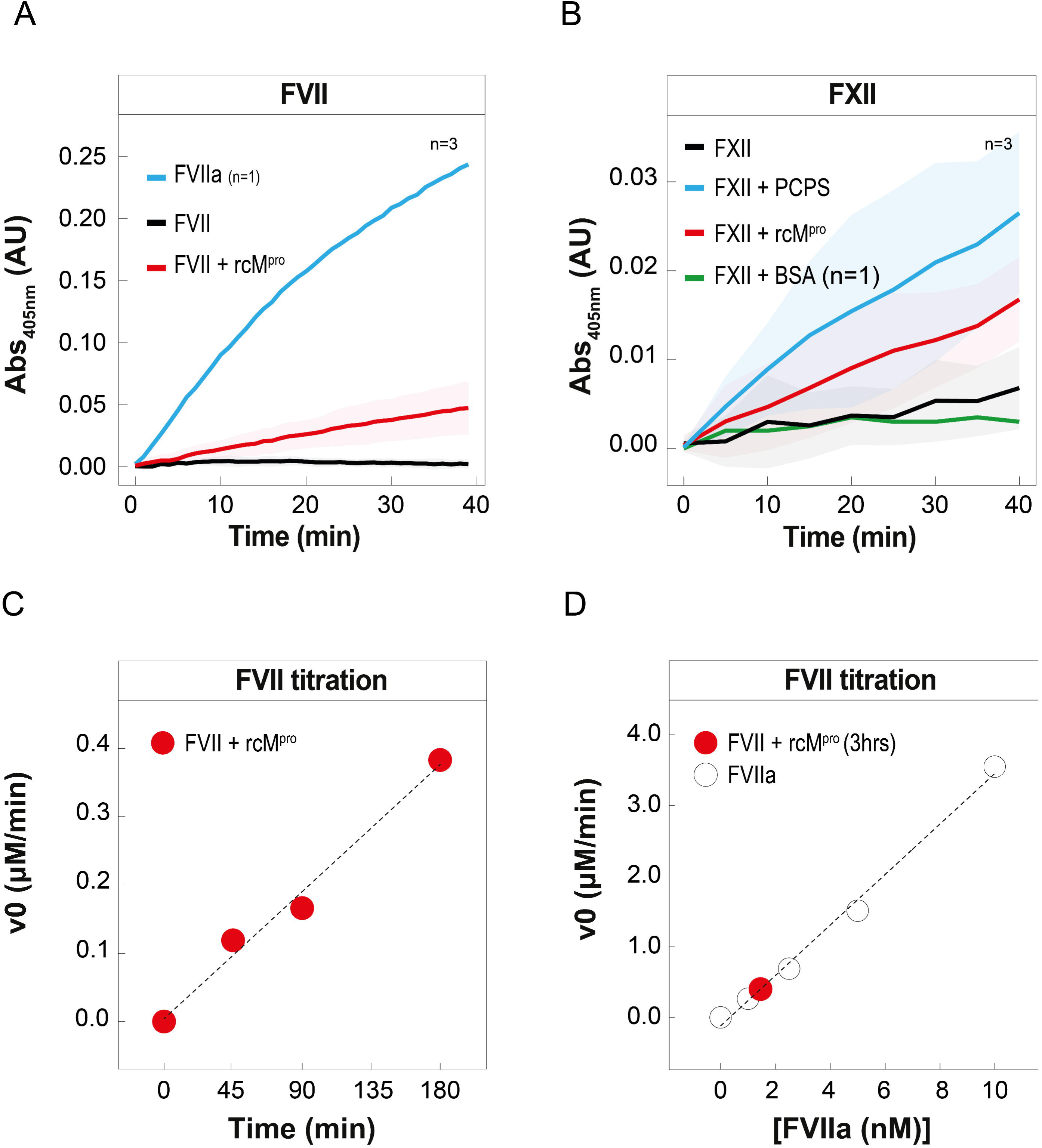
Analysis of the effect of rcM^pro^ on the activation of FVII and FXII zymogens. (A) Activation of FVII by rc-M^pro^ monitored by enzymatic assay. FVII zymogen (200 nM) was pre-incubated at 37°C with rcM^Pro^ (2 μM) in HBS-PEG buffer, pH 7.4, containing 3 mM CaCl_2_. After 3-h reaction, an aliquot (5 μl) of the proteolysis mixture was taken and added to a solution of the FVIIa specific chromogenic substrate MeSO_2_-D-CHA-But-Arg-pNA (300 μM) containing 100 nM recombinant tissue factor (TF). The time course of pNA release was monitored at 37°C by recording the absorbance change at 405 nm obtained with FVII zymogen after 3-h pre-incubation with (⁃) or without (⁃) rcM^pro^. The positive control (⁃) corresponds to the kinetics of substrate hydrolysis by fully active FVIIa added at a concentration (10 nM) identical to that of the FVII zymogen in the proteolysis reaction with rcM^pro^. Data are presented as mean ± standard deviation. (**B**) **Activation of FXII by rc-M^pro^ monitored by enzymatic assay.** FXII zymogen (50 nM) was incubated at 37°C with rcM^Pro^ (50 nM) in HBS-PEG buffer, pH 7.4, containing 5 mM CaCl_2_ and the chromogenic substrate S2302 (300 μM). The appearance FXIIa activity was detected by recording the time course release of pNA at 405 nm (⁃). The negative and positive control experiments were carried out by incubating FXII zymogen with a S2302 solution in the absence (⁃) or presence (⁃) of PCPS (40μM) liposomes. A further control experiment was performed by incubating FXII zymogen with a S2302 solution in the presence of BSA (50 nM) (⁃), as a negatively charged protein. Data are presented as mean ± standard deviation. (**C**) **Rate of FVIIa substrate hydrolysis at increasing pre-incubation times.** The initial rate (v_0_) of pNA release from the FVIIa-specific substrate was determined under the same experimental conditions as those reported in panel A, at increasing time of incubation of FVII zymogen with rcM^pro^ in the time range 0-3h. (**D**) **Quantitative determination of active FVIIa generated by rcM^pro^-catalysed cleavage of FVII zymogen**. The initial rate (v_0_) of substrate hydrolysis by active FVIIa is plotted as a function of increasing FVIIa concentrations (○). The calibration curve, resulting from the linear interpolation of the data points, was used to estimate the concentration/amount of active FIIa which is generated after 3-h incubation of FVII zymogen with rcM^pro.^(●).

The amount of newly generated FVIIa after 3-h reaction of FVII with rc-M^pro^, was estimated using a titration curve, obtained from the plot of the initial velocity (v_0_) of substrate hydrolysis as a function of FVIIa concentration (**Figure S4A,B**), and found to be approximately 14% of the total amount of FVII zymogen originally present in the activation reaction (**Figure 3C,D**). FXII is physiologically activated by binding to negative surfaces, that stabilise the zymogen in a catalytically competent conformation ^70^. To rule out the possibility that rcM^pro^, being negatively charged (pI= 5.9) at physiological pH conditions, could provide *per se* a negatively charged surface for the conformational activation of FXII, we compared the effect of rcM^pro^ to that observed with bovine serum albumin (BSA, pI = 5.9). The data in **Figure 3B** indicate that BSA does not activate FXII, further supporting the hypothesis that M^pro^ activates FXII *via* a proteolytic mechanism.

Overall, the results of this screening highlight FVII and FXII as potential substrates for rcM^pro^ in human plasma, whereby the two zymogens are strategically positioned at the beginning of the extrinsic and intrinsic pathway, respectively, of the coagulation cascade.

### 3.3. M^pro^ displays secondary substrate specificity for Arg-X peptide bonds

Proteolytic activation of coagulation factor zymogens is strictly conserved during evolution, from Fish, Amphibians, Reptiles, Birds and Mammals, and requires the cleavage at Arg-X bond, where X is a hydrophobic amino acid (usually Val or Ile)^25^. In humans, FVII is physiologically activated by FXa, FXIIa, FIXa, or thrombin (FII) by cleavage at Arg152-Ile153 bond^71^, while FXII is cleaved at Arg334-Asn335, Arg343-Leu344, and Arg353-Val354 bonds. However, only cleavage at Arg353 leads to the generation of αFXIIa active form^72,73^. Clearly, the requirements for activating these zymogens stand in stark contrast to the canonical cleavage specificity of M^pro^, that mandates the presence of a Gln at P1 and a hydrophobic residue (Leu/Met) at P2, resulting in the precise cleavage of 11 sites within viral polyproteins 1a and 1ab^41–43,74,75^.

Starting from these considerations, we decided to more deeply investigate M^pro^ substrate specificity by applying a modified version of the HTPS (High Throughput Protease Screen)^41^ workflow for cysteine proteases, as detailed elsewhere^54,55^. We applied this strategy to capture any potential secondary substrate specificity of M^pro^ by analysing the identified cleavages of the protease with time resolution (5 min – 24 hours). Briefly, we incubated rcM^pro^ with a native standardized cell extract (E:S 1:10) and identified by mass spectrometry (MS) the peptide fragments released by the protease (for details, see the Methods and legend to **Figure 4A**). These peptides were then used to generate a frequency/position amino acids matrix showing the preference of M^pro^ for amino acids from position P4 to P4’, according to the Schechter and Berger nomenclature^76^ (**Figure 4A**).

**Figure 4.**
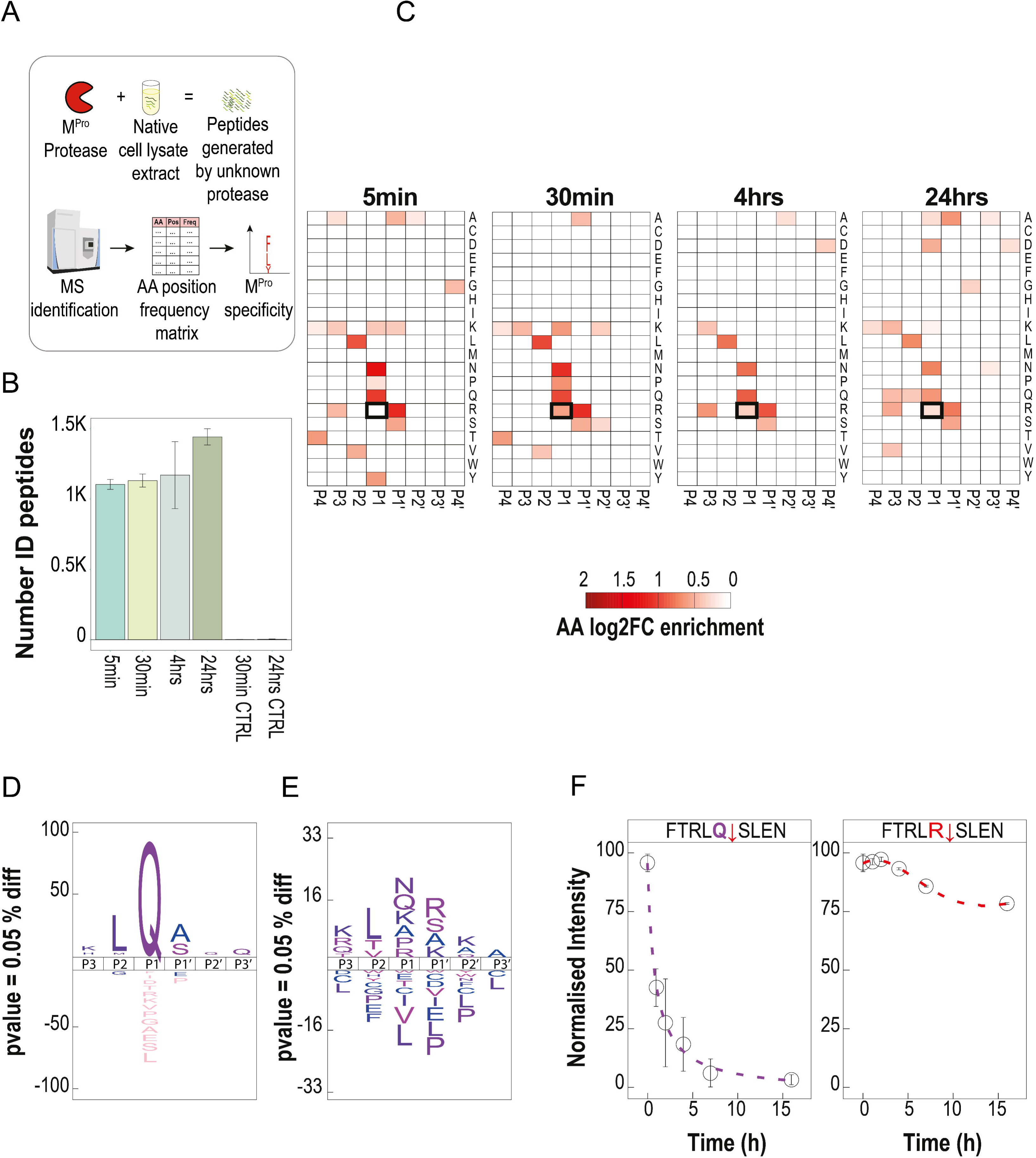
Characterization of M^pro^ substrate specificity. (A) HTPS workflow. Briefly, rcM^pro^ is incubated with a native standardized cellular extract at different time points. The peptide fragments, generated under native conditions, are isolated by filter-aided sample preparation (FASP) and subsequently identified by data dependent acquisition (DDA) mass spectrometry (see Methods). Significant enrichment (*p* value < 0.01) of amino acid frequency, compared to a random distribution, enables generating in triplicate an amino acid-position frequency matrix and the profiling of M^pro^ substrate specificity. (**B**) **Bar plot of identified peptides/cleavages in HTPS assay.** Peptides are generated incubating rcM^pro^ in the standardized cellular extract for 5min, 30min, 4hours and 24 hours. As a control, background peptides are identified in the absence of rcM^pro^ after 30-min and 24-hours incubation time (see Methods). Histograms refer to the average number of peptides generated in three independent assays (n=3) at each time point, with error bars corresponding to the standard deviation. (**C**) **Positional substrate preferences of M^pro^ (from P4 to P4’) determined by HTPS assay.** The heatmaps report the amino acid enrichment (log_2_FC) compared to random amino acid distribution. Significant (p value < 0.01) and positive enrichment are shown using a red color palette. Specificity of M^pro^ at each time point is calculated from three independent replicate experiments (n=3). The respective natural amino acids are sorted alphabetically. (**D/E**) **Substrate specificity of M^pro^.** IceLogo plot of substrate specificity for P3-P3’ positions of M^pro^ after 30-min reaction. The specificity is reported considering cleavages with Gln in P1 position (**D**) and for all identified cleavages (**E**). (**F**) **Time-course analysis of peptide substrate hydrolysis by rcM^pro^.** The synthetic peptide FTRL**Q**↓SLEN and its analogue FTRL**R**↓SLEN were separately incubated (10µM) in HBS at 37°C with rcM^pro^ (E:S molar ratio 1:40). The relative amount of residual intact peptides was quantified by LC-MS at increasing time points (**Figure S6**). Measurements were carried out in three independent experiments, with error bars corresponding to the standard deviation ±SD at each time point.

After incubating the native cell extract with rcM^pro^ for 5, 30min, 4 and 24hours, we identified 1105±31, 1139±42, 1173±238 and 1442±55 unique cleavages, respectively (**Figure 4B, Table S1**). The substrate preference, calculated as log_2_-fold change (log_2_FC) enrichment of amino acids compared to a random distribution, confirmed the canonical specificity for M^pro^, with Gln in P1, Leu and Val in P2, and Ser/Ala in P1’ position at all measured time points (**Figure 4C,D**). Notably, even after 5-min incubation, there is a significant (*p* value < 0.01) log_2_FC enrichment >1.0 for Leu in P2 and Gln at P1 position (**Figure 4C,E**). The analysis indicates also an enrichment of Arg at P1’ position, as well as Asn at P1 (**Figure 4C,E**). The latter result is not surprising, as Asn and the canonical Gln share the same polar carboxamide-group. The emerging preference for Arg at P1’ is in line with the observation that the S1’ site displays a negative electrostatic potential in M^pro^ structure (see section **3.4** and **Figure 5D**). The time resolution dimension in this analysis allows to probe the changes in M^pro^ substrate preferences. Only after 30-min incubation, rcM^pro^ displays an emerging secondary specificity for basic amino acid residues (Arg and Lys) at P1 position (with a log_2_FC = 0.6, p value < 0.01), which is maintained up to 24 hours only for Arg (**Figure 4C, Table S1**). During the HTPS analysis, we identified >10^3^ cleavage sites containing an Arg-residue in position P1, corresponding to about 10% of all the cleavage sites identified.

**Figure 5.**
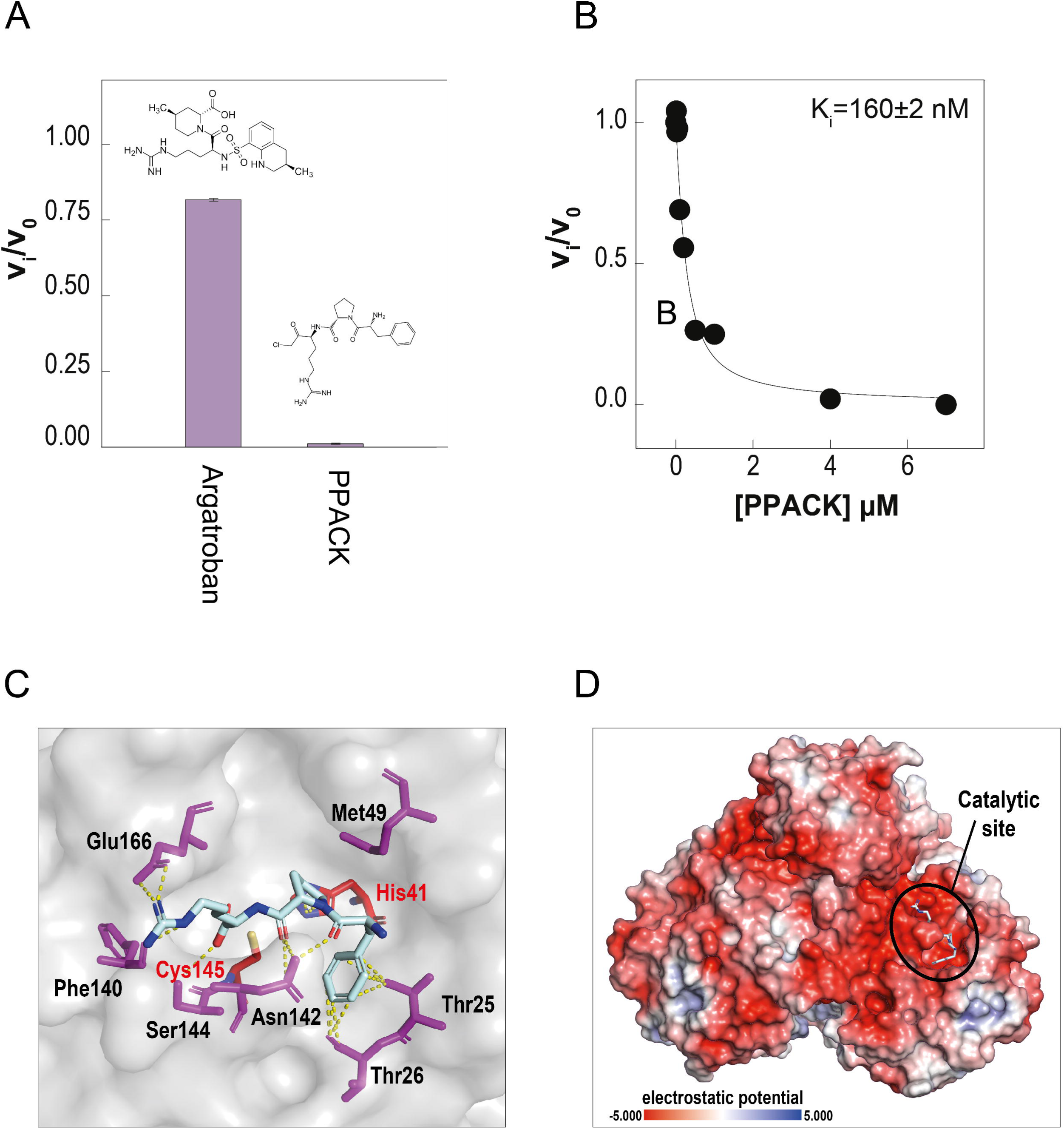
Probing PPACK-M^pro^ interaction by enzyme inhibition assays. (A) **Bar plot of M^pro^ inhibition by argatroban and PPACK.** Argatroban or PPACK (7μM) were incubated for 2 h in HBS-PEG, pH 7.4, at 37±0.1°C with rcM^pro^ (50 nM). The reaction was started by addition of the fluorogenic substrate (1.25μM) and the rate of hydrolysis was determined by recording the fluorescence increase at 530 nm. The data are expressed as the relative rate (v_i_/v_0_) of substrate hydrolysis in the presence (v_i_) and absence (v_0_) of the inhibitor. (B) **Determination of M^pro^ inhibition constant by PPACK.** Enzyme inhibition assays were carried out as in panel **A**. The values of (v_i_/v_0_) are plotted as a function of inhibitor concentration. The data points were interpolated with **equation 2**, describing the tight binding inhibition model, to extract the apparent inhibition constant (K_i_ ^app^), from which a value of K_i_ = 160±2 nM was estimated (see Methods). (**C**) ***In silico* docking of M^pro^ with D-Phe-Pro-Arg tripeptide.** Close-up view of D-Phe-Pro-Arg peptide (shown in stick) (i.e., the peptidyl moiety of PPACK D-Phe-Pro-Arg-CH_2_Cl) docked into M^pro^ active site (shown as van der Waals surface, light grey). Ligand atoms are colour coded: nitrogen in blue, oxygen in red, carbon in dark grey. Relevant amino acid side-chains in M^pro^ are indicated. The Arg guanidyl-group is electrostatically coupled to Glu166 in the S1 site, whereas D-Phe points toward His41 in the active site (AS) whereas Pro interacts with Thr25 and Thr26 in the S1’ sub-site. HPEPDOCK software was used in docking simulations (see Methods). (**D**) **Surface electrostatic potential of M^pro^ in the dimeric form**. Calculations were carried out using the APBS software on the crystallographic structure of M^pro^ in the dimeric form (6y2e.pdb). The surface is coloured according to the electrostatic potential (blue, positive; red, negative), as indicated, and expressed as kJ/(mol·q).

To rule out any possible contribution from trace amounts contamination of proteases in the native cell lysate, we performed HTPS experiments in the absence of M^pro^ (mock control) after 30-min and 24-hours incubation. Noteworthy, we identified less than 10 background peptides *per* condition, thus excluding any bias generated from active proteases present in the native standardized cell lysate (**Figure 4B** and **Table S1**). Importantly, no proteases with Arg specificity were used in the purification of rcM^pro^ and rM^pro^ that were utilized in this work. However, as an additional control, we checked whether trace amounts of proteases other than M^pro^, eventually present in preparations of rcM^pro^ samples, could affect the results of the HTPS screen. SDS-PAGE and intact mass analysis confirm the purity of the sample and the presence of rcM^pro^ in the monomeric and dimeric state (**Figure S5A,B**). Furthermore, using bottom-up proteomics analysis of rcM^pro^ preparations treated with two different proteases (i.e., Trypsin and Glu-C specific endoproteinase from *S. aureus*) and searching the resulting spectra against an extended database (Uniprot pan proteomes – https://ftp.uniprot.org/pub/databases/uniprot/current_release/knowledgebase/pan_proteomes/), at the best of the sensitivity of our MS analysis, we could not identify any protease sequence in the rcM^pro^ preparations tested (**Table S2**).

Similar to our findings, previous proteomic studies (i.e., N-terminome analyses) indicated that M^pro^ exhibits an extended specificity with the possibility to cleave after a Met-or His-residue at P1 position^30^. Despite HTPS data cannot confirm this additional specificity for M^pro^, the possibility that an Arg-residues can be found in the P1 substrate specificity site confirm the broader specificity of M^pro^ previously highlighted and is fully consistent with the observed proteolytic activation of FVII and FXII zymogens by rcM^pro^, emerging from our enzymatic assays (**Figure 3**).

To validate HTPS findings on the secondary specificity of M^pro^ for Arg-X bonds and further corroborate the hypothesis that the protease can play a role in the pathological activation of blood coagulation in SARS-CoV-2 infections, based on the canonical substrate preference of M^pro^ ^77^, we synthesised a putative peptidyl substrate FTRL**Q**↓**S**LEN and its analogue FTRL**R**↓**S**LEN in which the canonical Gln in P1 position was replaced with an Arg-residue. The data shown in **Figure 4F** and **Figure S6** indicate that, although the efficiency at which the analogue is hydrolysed by the protease is much lower, the presence of an Arg-residue in P1 position, replacing Gln, does not inhibit the cleavage by rcM^pro^.

Overall, the integration of HTPS screening dataset with the biochemical characterization of synthetic peptidyl substrates put forward the possibility that M^pro^ has an extended substrate specificity and can proteolytically activate both FVII and FXII to trigger plasma clotting.

### 3.4 M^pro^ can accommodate an Arg-residue in the substrate primary specificity site

In the previous chapter, we have shown that M^pro^ displays an extended/secondary substrate specificity allowing cleavage of substrates with an Arg at P1, resembling in some way the substrate specificity of coagulative proteases^25,54^. Here, we provide biochemical and structural data on the mechanism underlying the interaction of Arg-containing substrates/inhibitors with the protease active site.

Even though M^pro^ and α-thrombin belong to different protease families (i.e., α –thrombin is a serine protease, whereas M^pro^ is a cysteine protease), they share a common α-chymotrypsin fold^39,78^ with M^pro^ active site (i.e., His41 and Cys145) positioned at the interface of two orthogonal β-barrels (i.e. domains I and II). At variance with α-thrombin, an additional globular cluster of five helices (i.e., domain III) regulates protein dimerization in M^pro^ structure^40^. The structural similarity highlighted above for α-thrombin and M^pro^ can be also suggestive of similar molecular recognition properties of the two proteases toward both substrates and inhibitors. Hence, we challenged the viral protease with two highly specific α-thrombin inhibitors, i.e. argatroban and PPACK (D-Phe-Pro-Arg-CH_2_Cl) (**Figure 5A**). Argatroban is a reversible thrombin inhibitor (K_i_ = 19 nM), whereas PPACK irreversibly inhibits the enzyme (K_i_ = 24 nM) by forming a covalent bond with the catalytic His57 side chain. Noteworthy, both inhibitors orient an Arg-side chain in the protease S1 sub-site *via* electrostatic coupling with Asp189^79^. The results of rcM^pro^ inhibition assays indicate that, after 2-h incubation with excess inhibitor (1:140 E:I molar ratio), the rate of substrate hydrolysis by rcM^pro^ is reduced by approximately 20% with Argatroban while PPACK totally abrogates the catalytic activity (**Figure 5A**). Systematic analysis of rcM^pro^ inhibition by PPACK yields a K_i_ = 160±2 nM (**Figure 5B**), comparable to that earlier reported for thrombin ^27,79^.

Next, we mapped the PPACK binding region on M^pro^ structure by Hydrogen-Deuterium Exchange Mass Spectrometry (HDX-MS), an emerging technique in structural biology useful for investigating protein conformation, dynamics and molecular recognition^57,80–82^. Briefly, HDX-MS exploits the intrinsic propensity of backbone amide hydrogens at exposed/flexible sites to exchange more rapidly with deuterium than those hydrogens that are buried in the protein interior or at the ligand-protein interface and therefore these will exchange much more slowly. HDX can be monitored by recording the time-dependent mass increase of peptides, generated after proteolysis with pepsin, allowing to study protein dynamics and ligand-protein interaction with a spatial resolution of 3-5 amino acids and, importantly, with tiny sample amount (20-50 μg)^81^. HDX-MS analysis of rcM^pro^ was performed at 20°C in PBS, pH 7.4, before and after addition of PPACK. A coverage of ∼80% in the M^pro^ sequence was obtained, allowing us to estimate the extent of deuterium uptake (ΔHDX) for each peptide at increasing incubation times, from 15s to 2h (**Figure 6A, Table S3**). To identify the regions that are conformationally altered upon inhibitor binding, a difference three-dimensional map of deuterium uptake, after 2-h incubation with D_2_O, was generated on the crystallographic structure of M^pro^ in the dimeric active form (6y2e.pdb)^83^ (**Figure 6B**). HDX-MS data indicate that inhibitor binding induces shielding/ordering of well-defined regions in M^pro^ (**Figure 6A**). In particular, the segment 38-44 undergoes a significant decrease (i.e., –0.7 Da) in deuterium uptake upon PPACK binding, along with residues 88-90 which are located in the adjacent β-strand. Notably, peptide 38-44 comprises the catalytic His41. Furthermore, fragments 141-150 and 142-150, both encompassing the catalytic Cys145, could be identified solely in the PPACK-unbound form of the enzyme, as in the bound form they are covalently derivatized (**Figure S7B**). Altogether, these observations indicate that the inhibitor covers the active site and covalently binds to Cys145. The results of HDX-MS analysis are consistent with molecular docking simulations (**Figure 5C**) and electrostatic calculations (**Figure 5D**), showing that the positively charged Arg-side chain of PPACK harbours the negative Glu166 in the protease S1 site, whereas Pro points toward His41 in the active site and D-Phe contacts Thr25 and Thr26 in the S1’ sub-site on M^pro^ structure. As expected from its acidic pI value, the surface electrostatic potential of M^pro^ dimer is largely negative, in keeping with the observation that, among the annotated interactors of M^pro^, there are numerous basic proteins, i.e. histones and histone-related proteins, ribosomal and ribonuclear proteins ^38^.

**Figure 6.**
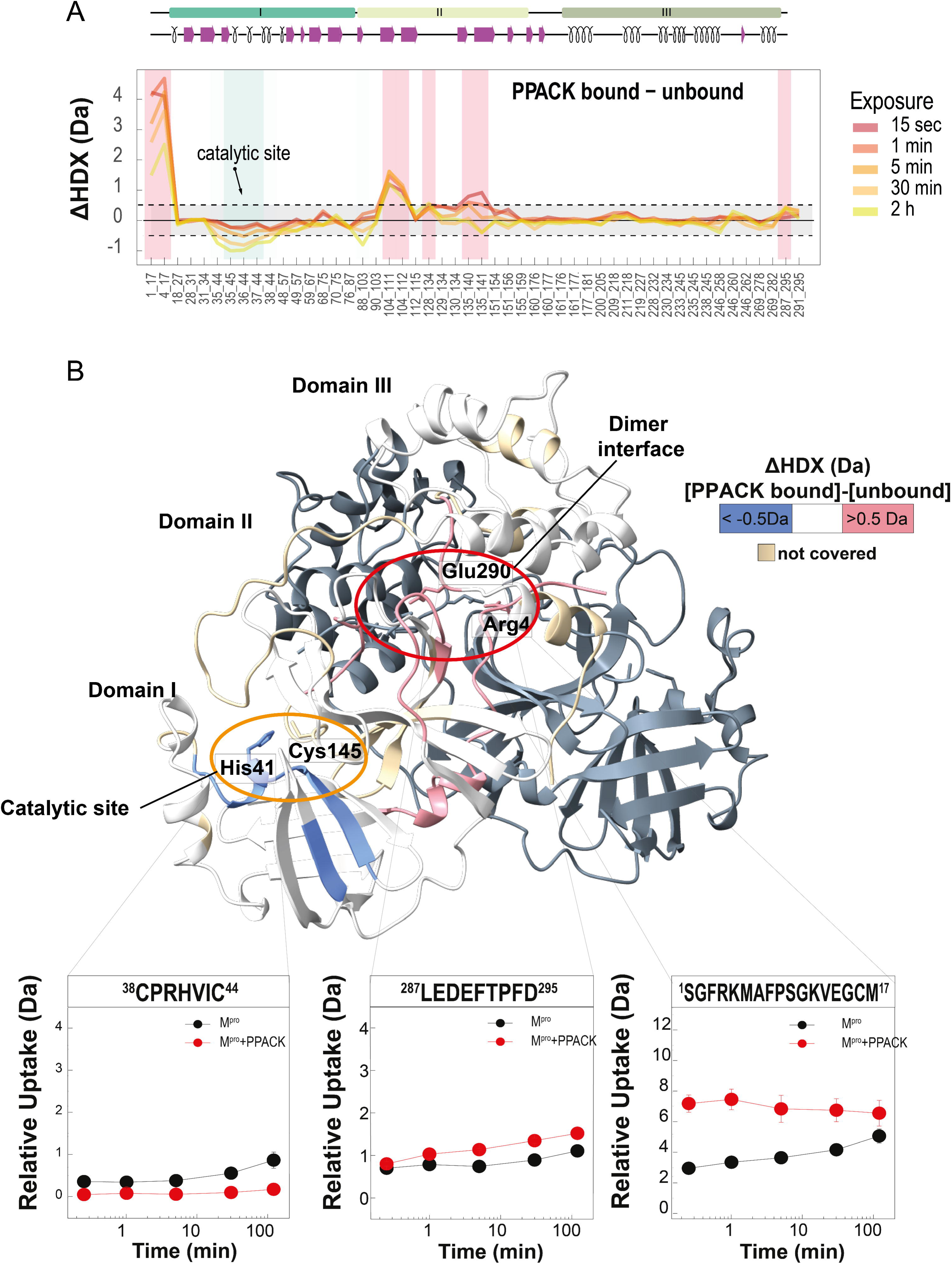
Probing PPACK-M^pro^ interaction by HDX-MS analysis. (**A**) **HDX-MS analysis of M^pro^-PPACK interaction**. The difference deuterium uptake (ΔHDX) profile, measured for peptide fragments deriving from rcM^pro^ in the PPACK-bound and –unbound form, is reported in the time range of D_2_O labelling 15s – 2h, as indicated (see Methods). Negative values of ΔHDX correspond to regions that become more shielded/rigid upon inhibitor binding, whereas positive values correspond to regions that become more exposed/flexible. The active site (yellow circle) and the dimerization region (red circle) are indicated. Experimental conditions were as follows: rcM^pro^ (1.25 μM) incubated at 20 °C in 20 mM PBS in 95:5 D_2_O:H_2_O solution, pD 7.4, containing 150 mM NaCl. At each time point, HDX-MS measurements were conducted in triplicate, with error bars as ±SD. (**B**) **Three-dimensional difference map of M^pro^ deuterium uptake after PPACK binding.** The values of ΔHDX, as obtained from HDX-MS data of rcM^pro^ in the PPACK-bound and –unbound state after 2-h exchange (panel **A**), are projected onto protomer A in the dimeric M^pro^ structure (6y2e), represented as a ribbon drawing. The regions that are more (ΔHDX < –0.5 Da) or less (ΔHDX > 0.5 Da) protected from H/D exchange after PPACK binding are coloured blue or red; the regions for which ΔHDX values are not statistically significant are in light grey, whereas those regions that could not be covered in HDX analysis are coloured gold. For the sake of clarity, protomer B is in grey. The relative deuterium uptake plots of relevant peptides in the active-site region (sequence 38-44), and at the N-terminal (sequence 1-17) and C-terminal (sequence 287-295) regions, in the PPACK-bound and –unbound states, are also reported.

Furthermore, HDX-MS data indicate that, besides shielding/ordering of the active-site cleft, rather surprisingly, PPACK binding induces marked exposure/flexibilization of the N-terminal (sequence 1-17) and C-terminal (sequence 288-295) regions comprising key residues (i.e., Arg4 and Glu290) forming inter-monomer salt bridges which are crucial for protease dimerization and associated function^83^. Likely, covalent PPACK binding destabilizes long-range these interactions, favouring dimer dissociation, that result in the exposure of the N– and C-terminal regions and final protease inactivation.

The results of enzyme inhibition assays and HDX-MS analysis with PPACK provide structural support to the hypothesis that M^pro^ can allocate an Arg-side chain in the S1 sub-site, thus explaining on structural grounds the secondary substrate specificity of the protease for Arg at P1, emerging from HTPS analysis (**Figure 4**).

### 3.5 Identification of the cleavage sites for rcM^pro^ on coagulation factors VII and XII

The discovery of M^pro^ secondary preference for Arg at P1 position suggests that M^pro^-mediated activation of FVII and FXII might occur via canonical proteolysis at Arg-residues, as earlier reported for known physiological activators ^71,73^. To verify our hypothesis, we identified the M^pro^ activation cleavage sites on purified FVII and FXII by exploiting a simplified version of the Terminal Amine Isotopic Labelling of Substrates (TAILS) protocol^54,60^. This method allows to identify new N-termini on a given protein, possibly generated upon proteolysis, by reductive di-methylation reaction followed by trypsin cleavage and classical bottom-up MS workflow. The differential analysis of the N-terminally dimethylated peptides from the M^pro^-treated and untreated samples (mock control) allows identification of the new N-termini, corresponding to the cleavage sites (**Figure 7A**).

**Figure 7.**
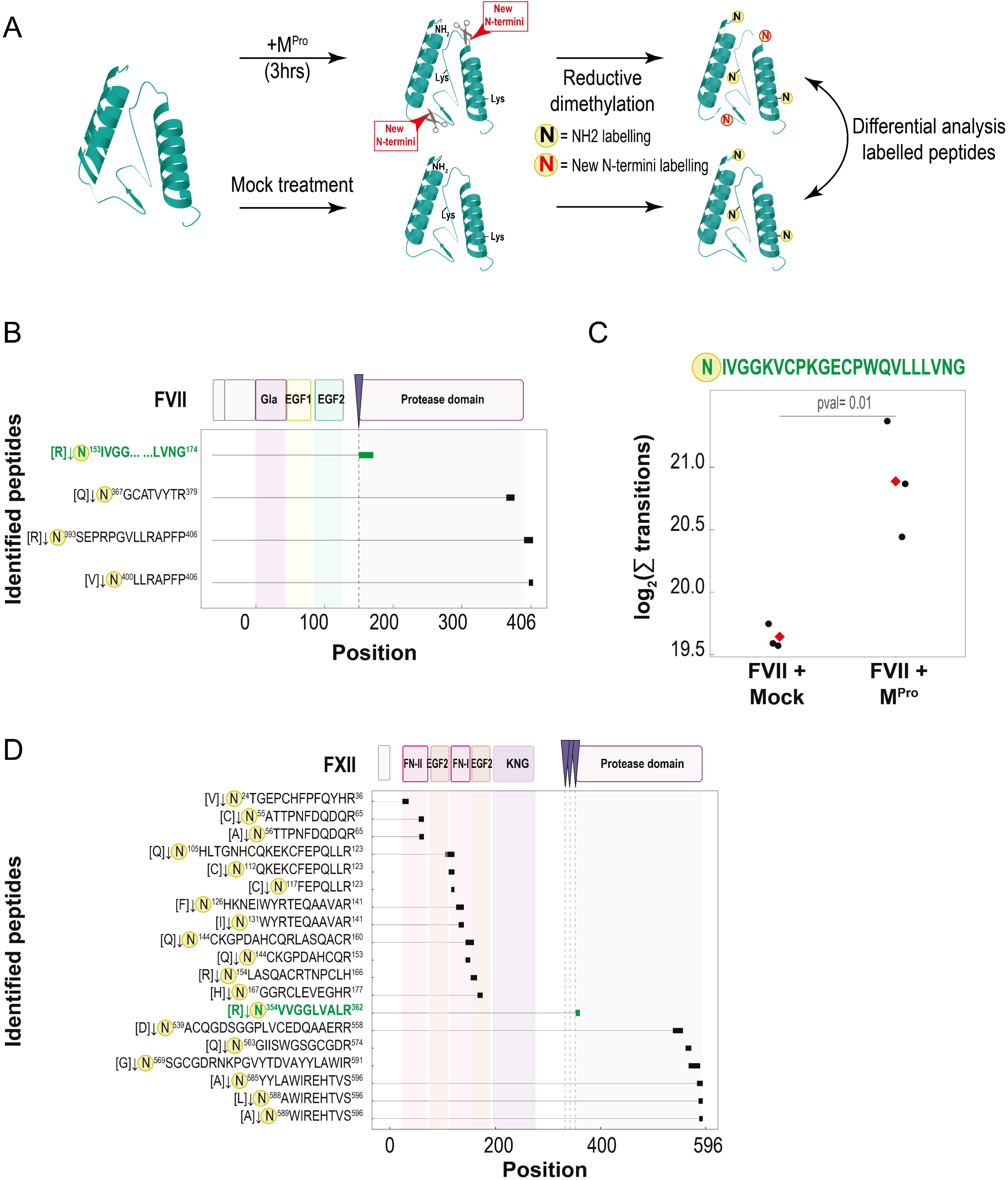
Identification of rcM^pro^ cleavages in FVII and FXII zymogens by TAILS analysis. (**A**) **TAILS workflow.** Simplified version of TAILS workflow in which the substrates (FVII and FXII) are proteolyzed with M^pro^ (1:10 E:S molar ratio) and mock treatment. Generated cleavage fragments are subsequently di-methylated under reducing conditions. Di-methylated peptides are identified by mass spectrometry and subjected to differential mass analysis to identify M^pro^ cleavage sites. (**B**) **M^pro^-specific cleavages mapped in the FVII sequence.** Results from TAILS experiment using rcM^pro^ and mock treatment. The experiment is performed with three independent replicates. Cleavages identified in at least two replicates (except ^153^IVGGKVCPKGECPWQVLLLVNG^174^) and not in the mock control are shown. Dashed lines indicate the cleavage site (Arg152-Ile153) for FVII physiological activators. The peptide ^153^IVGGKVCPKGECPWQVLLLVNG^174^ generated by the cleavage in the activation site is annotated in red. (**C**) **Quantitative determination of** ^153^**[DimethNter]IVGGKVCPK[DimethLys]GECPWQVLLLVNG**^174^ **peptide.** The peptide is quantified using a targeted mass spectrometry approach in both the mock control (FVII alone) and in the presence of rcM^pro^. The intensity reported is calculated from the sum of the top five peptide transitions. This experiment was conducted with three replicates, with each circle denoting a single measurement and the diamond indicating the mean value. (**D**) **M^pro^-specific cleavages mapped in the FXII sequence.** Results from TAILS experiment using rcM^pro^ and mock treatment. The experiment is performed with three independent replicates. Cleavages identified in at least two replicates and not in the mock controls (t = 0 and 3h) are shown. Dashed lines indicate the cleavage sites for FXII physiological activation.

After 3-h incubation of FVII with rcM^pro^, we identified three N-terminal di-methylated peptides in at least two over three biological replicates, resulting from the cleavage at Gln366-Gly367, 392Arg-Ser393 and Val399-Leu400 bonds (**Figure 7B** and **Table S4**). The cleavages were absent in the mock control. The identification of cleavage sites different from the canonical Gln at P1 position, is consistent with the extended substrate specificity highlighted above for M^pro^ by HTPS analysis (**Figure 4C**). All three cleavages are located at the fraying C-terminal region of FVII zymogen protease domain and likely not involved in zymogen activation. Furthermore, in one of the three replicates (but not in the mock control), we could identify the cleavage of rcM^pro^ at the Arg152-Ile153 bond (**Figure 7B**, green peptide in narrow), matching with the canonical activation site on FVII zymogen by its physiological activators, i.e. FXa, FXIIa, FIXa, and thrombin^71^. As the generation of ^153^IVGGKVCPKGECPWQVLLLVNG^184^ can be taken as a safe marker of FVII zymogen activation, we leveraged the Parallel Reaction Monitoring (PRM) method^84^ to increase sensitivity and selectively measure the intensity of the transitions of the aforementioned peptide (**Figure 7C, Table S4**). The quantitative analysis revealed that, after 3-h incubation with rcM^pro^, there are residual trace amounts of the activation peptide even in the mock control samples, consistent with the autoactivation of the FVII zymogen earlier reported^85^. Furthermore, we found a significant (*p-*value = 0.001, unpaired two-sided Student’s t-test) two-fold increase in the intensity of the activation peptide in rcM^pro^-treated samples, compared to controls. Even though unspecific proteolysis at the very C-terminal region in FVII might in theory lead to a loss of function in the resulting truncated species, the generation of active FVIIa in the proteolysis reaction of FVII zymogen with rcM^pro^, as experimentally observed in **Figure 3A-C**, indicate that the cleavage at the physiological activation site (i.e. Arg152-Ile153) is prevalent.

The same TAILS strategy, highlighted above for FVII, was exploited to identify the cleavage sites for rcM^pro^ on FXII zymogen. After binding to negatively charged membranes, FXII zymogen is cleaved *in vivo* primarily at Arg334-Asn335, to generate active FXIIa protease, and eventually at two other two extra bonds, i.e. Arg343-Leu344, and Arg353-Val354. After incubation of FXII zymogen in solution with rcM^pro^ (in the absence of negative liposomes), TAILS analysis allowed us to safely identify 19 N-terminal di-methylated peptides in at least two out of three replicates, that are absent in the mock controls (t = 0 or 3 h) (**Figure 7D**). The variable nature of the amino acids at P1 position in the cleaved peptide bonds once more supports the extended specificity of M^pro^, highlighted in this study (**Figure 4C**). Most of the cleavage sites that are mapped in **Figure 7D** are localized in the N– and the C-terminal region of the zymogen, outside the FXII protease domain and therefore not relevant to zymogen activation. Importantly, our TAILS analysis identifies in all three replicates but not in the control the peptide ^354^VVGGLVALR^362^, generated from the cleavage at the physiological activation site, i.e. Arg353-Val354 bond. The latter finding clearly indicates that rcM^pro^ can proteolytically activate FXII zymogen.

Overall, applying the TAILS workflow and the PRM method on purified protein zymogens, we dug into the activation mechanism of FVII and FXII by M^pro^ and discovered that the protease can directly activate FVII and FXII by proteolysis at their physiological activation sites, i.e. Arg152-Ile153 and Arg353-Val354 bond, respectively, fully consistent with the results of enzymatic assays (**Figure 3**).

## 4. DISCUSSION

Thrombotic diseases may exist as a primary/autonomous (i.e., idiopathic) pathological state or appear as the expression of secondary complications, occurring with variable incidence and severity, in different (apparently unrelated) diseases, having different etiology and clinical phenotype, such as for instance type-2 diabetes (T2D) ^86^, chronic kidney disease (CKD) ^87^, inflammatory bowel disease (IBD) ^88^, cancer ^89^, rheumatoid arthritis ^90^ autoimmune diseases ^91^

^92^ ^93^, amyloid diseases ^94^ ^95^ ^96^ ^97^ ^98^, and bacterial ^99^ ^100^ and viral ^101^ infections. Even though in most cases the molecular understading of secondary thrombotic complications remains elusive, non-canonical mechanisms leading to the activation of blood coagulation have been proposed, including oxidative modifications of von Willebrand factor in T2D ^102^ ^103^ and CKD ^104^ or aberrant proteolytic activation of coagulation factors in IBD ^105^.

The link between COVID-19 and a markedly increased risk of thrombosis has been clear since the early days of SARS-CoV-2 pandemic, with COVID-19-associated coagulopathy strongly contributing to morbidity and mortality in infected patients^106^. Several evidence indicate that COVID-19 coagulopathy is related to the innate immune response after virus infection. These include the overproduction of pro-inflammatory cytokines “cytokine storm”^20,21^ that lead to the increased expression of Tissue Factor and to the formation of neutrophil extracellular traps that promote platelet activation ^107^. This phenomenon is further amplified by the “bradykinin storm”, a positive feedback loop in which FXIIa, besides activating FXI, triggers the kallikrein-kinin system, leading to the release of bradykinin (BK) and other inflammatory cytokines^108^. During the SARS-CoV-2 infection, the viral protease M^pro^ proteolytically process multiple host proteins, modifying the proteome composition and enforcing virus infection and maturation^30–32,109^. Two lines of evidence indicate that M^pro^ can circulate in the blood: i) the presence of anti-M^pro^ antibodies in the sera of COVID-19 patients^33,34^ and ii) the identification of SARS-CoV-2 mRNA in the bloodstream and in brain tissue ^35,110^.

Building on our previous discovery that the extracellular serine protease subtilisin from *Bacillus subtilis*, a non-pathogenic Gram-positive bacterium in the human gut microbiome, can directly activate the coagulation cascade^59^, this study investigates whether the M^pro^ protease secreted by the SARS-CoV-2 virus, circulating in the bloodstream, can similarly activate coagulation factors. This could provide another causal relationship for the positive correlation between a pro-thrombotic state and COVID-19 severity ^106^. To assess whether M^pro^ can trigger plasma clotting, we tested different protease preparations (rcM^pro^ and rM^pro^) and both fresh and commercial reconstituted plasma samples. Turbidimetric assays show that the addition of M^pro^ to human plasma is a sufficient condition to promote coagulation in all the protease-matrices combinations, representing the first evidence that M^pro^ exhibits an intrinsic procoagulant activity. This effect was further confirmed with plasma samples from a cohort of 18 healthy donors, which displayed higher clotting probability in the presence of M^pro^ compared to the control group (p-value = 0.022). Having established that M^pro^ can induce clot formation, we asked whether the protease could directly activate specific coagulation factors in vitro (e.g., M^pro^ added to the zymogen solution containing the corresponding chromogenic substrate) and found that, among the zymogens tested, M^pro^ solely activates FVII and FXII, which are strategically positioned at the beginning of the extrinsic and intrinsic pathway of blood coagulation, respectively. The results of TAILS analysis clearly indicate that M^pro^ cleaves FVII at multiple sites and in particular at the Arg152-Ile153 bond, which is the physiological activation site of FVII by FXa, FXIIa and FIXa^71^. Notably, this cleavage results in the activation of approximately 14% of treated FVII. Likewise, FXII zymogen is cleaved by M^pro^ at multiple sites in the heavy chain and, importantly, at the physiological activation site Arg353-Val354, which may become exposed after proteolysis of the heavy chain. Altogether, these findings disclose a novel (i.e., proteolytic) mechanism that, in addition to the “cytokine and bradykinin storm” hypothesis ^11,20,21^, can contribute to explain at the molecular level the onset of thrombotic complications in severe SARS-CoV2 infections.

FVII circulates in the plasma at relatively low concentrations (0.5μg/ml), compared to other coagulation proteases, and approximately 1% of the zymogen is present in the active cleaved form (FVIIa). After proteolysis at the activation site, FVIIa retains a zymogen-like form which is converted to the fully catalytically competent conformation only after binding to TF ^68^. FXII circulates in the plasma in the closed form (30–40 μg/ml), with the activation site being shielded by the zymogen heavy chain. Upon interaction with negatively charged surfaces (e.g., the damaged endothelium), the heavy chain is displaced while the activation site becomes exposed, allowing (auto)proteolytic activation by cleaved/active FXIIa molecules present in solution or bound to the surface and by conformationally activated surface-bound FXII molecules ^111^. The ensuing FXIIa, besides activating FXI downstream in the coagulation cascade, can also convert prekallikrein to active kallikrein, which reciprocally activates FXII by proteolysis, thus amplifying FXIIa formation ^72^. Active kallikrein then stimulates the release of BK which in turn boosts downstream inflammation ^108^, typical of COVID-19 thrombotic complications ^24^. Once pathological activation of the coagulation cascade is initiated by M^pro^-mediated activation of FVII and FXII *in vivo*, the response could be amplified and propagated by multiple positive feedback reactions in the haemostatic network. As consequence, the generation of even tiny amounts of FVII and FXII are sufficient to induce final thrombin generation and thrombus formation ^68,112–114^. Therefore, we propose that M^pro^ could play a role in the development of the hypercoagulable state observed in SARS-CoV-2 infections. However, our data also indicate that the pro-thrombotic potential of M^pro^ displays significant inter-individual variability, ranging from negligible (if any) effect to a remarkable reduction in t_c_ values that has been observed in the clotting curves after addition of M^pro^ to plasma samples from healthy donors. This large variability observed *in vitro* is in keeping with clinical evidence accumulated so far showing that thrombotic complications manifest in only a limited number of COVID-19 patients^11^. Furthermore, these coagulative disorders are characterized by a wide spectrum of severity, ranging from sub-clinical hypercoagulable states to dramatic and even fatal thromboembolic events^11^. In this study, we did not systematically investigate the molecular determinants of the variability of the effect of M^pro^. However, literature data indicate that natural plasma levels of FVII and FXII remarkably vary within individuals, being affected by age, sex, genetic and environmental factors ^68,115^. Hence, we speculate that the lower susceptibility of some plasma samples to M^pro^, might be caused by the lower concentrations of FVII and FXII, naturally occurring in those samples. Although this hypothesis is reasonable, direct experimental verification is awaited in future work.

Cleavage of FVII and FXII at the physiological Arg-X activation sites could not be anticipated by the reported substrate specificity of M^pro^, dominated by the preference for a Gln-residue at P1 and a hydrophobic amino acid in P2 (Leu, Phe, Met) ^41–43,74,75^. A major achievement of this study is the identification by time-dependent HTPS profiling of an extended/secondary specificity of M^pro^, whereby the protease in addition to its canonical specificity (LQ↓) can cleave peptide bonds with Arg at P1, thus resembling the substrate specificity of coagulation factors ^54^. This finding is also reminiscent of recent degradomics studies, showing that M^pro^ can cleave substrates with His and Met at position P1 ^30,31^. More importantly, the previously undocumented substrate-specificity of M^pro^ for Arg-residues well explains at the molecular level our finding that M^pro^ can proteolytically activate FVII and FXII and trigger plasma clotting. The secondary substrate specificity of M^pro^ was further validated experimentally by i) enzyme inhibition assays, ii) HDX-MS analysis of inhibitor binding, and iii) enzymatic assays with synthetic peptide substrates. Starting from the structural similarity of M^pro^ and thrombin, we found that M^pro^ is efficiently inhibited by Arg-containing thrombin inhibitors argatroban and PPACK, orienting their Arg side-chain into the negatively charged S1 site of M^pro^, as demonstrated by HDX-MS analysis and supported by modelling studies. A conclusive proof that M^pro^ can cleave Arg-X bonds comes from the observation that the protease hydrolyses the peptide FTRL**R**↓SLEN, containing the non-canonical Arg at P1, albeit less efficiently than the canonical peptide substrate FTRL**Q**↓SLNE.

In conclusion, in this work we provided several pieces of experimental evidence showing that M^pro^ can induce plasma clotting by proteolytically activating FVII and FXII, which in turn can initiate the extrinsic and intrinsic pathways of blood coagulation, with a final pro-coagulant effect. This non-canonical mechanism highlights a possible novel function of M^pro^ *in vivo* that, in addition to the “cytokine and bradykinin storm” mechanism, can contribute to the pathogenicity of SARS-CoV-2 in COVID-19.

## Author contributions

VDF conceived and coordinated the work; AP, GN, API and LA performed the biochemical work; FU and AP performed proteomic and mass spectrometry analyses; MB, MLM, and BG provided in-house produced recombinant M^pro^; GDS, PM and PS provided plasma samples; PP supervised the TAILS tasks; VDF, AP, and FU designed research and jointly wrote the manuscript; all authors analyzed and interpreted the data, reviewed and approved the final content of the manuscript.

## Additional Information

### Funding

This work was supported by a Grant from the CaRiPaRo Foundation Excellence Research Project – BPiTA n. 52012 to VDF and a Grant from the Italian Ministry of University and Research (MIUR) – PRIN-2022 n. 2022ZSA2JP to PS and VDF.

### Conflict of interest

The authors declare that they have no conflicts of interest with the contents of this article.

## Supplementary Tables

**Table S1:** Detection of temporal dependent cleavages of M^pro^ by HTPS workflow and identification of substrate specificity.

**Table S2:** Identification of potential contaminants in the rcM^pro^ preparation. After digestion with Trypsin and GluC, peptides were search against the Uniprot pan proteomes.

**Table S3:** Characterization of the binding of PPACK with M^pro^ by HDX. Peptic peptides identified at different time in deuterium buffer as well the relative uptake difference are shown

**Tabls S4:** Identification of FVII and FXII cleavages upon incubation with M^pro^. The data includes both untargeted analysis (FVII and FXII) and targeted analysis (FVII only) of peptides generated using the TAILS workflow.

## Additional Information

### Funding

This work was supported by a Grant from the CaRiPaRo Foundation Excellence Research Project – BPiTA n. 52012 to VDF and a Grant from the Italian Ministry of University and Research (MIUR) – PRIN-2022 n. 2022ZSA2JP to PS and VDF.

### Conflict of interest

The authors declare that they have no conflicts of interest with the contents of this article.

## Supplementary figure legends

**Figure S1**. **Effect of M^pro^ on fibrin generation in human plasma.** (**A-C**) Clotting curves obtained after adding rM^pro^ (**A,C**) or rcM^pro^ (**B**) to human plasma samples collected from healthy donors (**A,B**) or reconstituted commercial plasma (**C**). Plasma samples were diluted 1:2 with HBS, re-calcified, and the increase of turbidity at 350 nm was measured over time at 37±0.1°C. Measurements were carried out in triplicate on a single plasma sample with shaded areas corresponding to the standard deviation ±SD at each time point. (**D**) Electrophoretic analysis of plasma clotting reaction. After 1-h incubation at 37°C of 1:2 diluted plasma with rM^pro^ (200 nM), the clot was collected, washed with HBS buffer and resuspended with Gdn-HCl 8M (incubated for 1h at 37°C). Albumin was then removed by treatment with EtOH to a final concentration of 42% for 1h at 4°C and following centrifugation at 16,000g for 45min at 4°C. After removing the supernatant, the pellet was resuspended with SDS sample loading buffer and analysed by reducing SDS-PAGE (Bolt Bis-TRIS 4-12% precast gel) and Coomassie staining (Simply Blue SafeStain, Invitrogen). For comparison, commercial fibrinogen (Fbg) was also loaded. The typical α-, β– and γ-chain of Fbg are indicated by arrows, along with the γ-chain dimer.

**Figure S2**. **Turbidimetric analysis of the effect of M^pro^ on fibrin generation in human plasma samples from healthy subjects.** Human plasma samples were collected from healthy donors, diluted 1:2 with HBS, and the increase of turbidity at 350 nm was measured over time at 37±0.1°C in the absence and presence of rcM^pro^, at the indicated concentrations. The curves are shown with the same scale (**Figure S2A**) and with a scale normalized for the maximum absorbance changes (**Figure S2B**) to facilitate the identification of sigmoidal profiles.

**Figure S3. Screening for M^pro^ ability to activate coagulation factors. (A) Activation of fibrinogen monitored by turbidimetric assay.** Clotting curves (n=2) obtained by adding α-thrombin (2nM) (⁃) or rcM^pro^ (100 nM) (⁃) to isolated fibrinogen solution (0.15mg/ml) in the presence of 1.5 mM CaCl_2_. (**B) Activation of prothrombin (FII) monitored by enzymatic assay.** Activation curve generated by S2238 (α-thrombin specific chromogenic substrate) (n=3). Prothrombin activity with rcM^pro^ (10:1 mol/mol E:S ratio) or without protease after 3-h incubation are shown as red and black curves, respectively. As positive control the activity of prothrombin is monitored after treatment with 1nM FXa in the presence of 100μM PCPS (50:50) (⁃, n=1). (**C**) **Activation of FX monitored by enzymatic assay**. Activation curve generated by S2765 (FXa specific chromogenic substrate) (n=2). FX activity with rcM^pro^ (10:1 (mol/mol) E:S ratio) or without protease after 3-h incubation are shown as red and black curves, respectively. As a positive control, the activity of FX is monitored after treatment with 0.2nM FVIIa in the presence of 2nM recombinant tissue factor (TF) (⁃, n=1). (**D**) **Activation of FIX monitored by enzymatic assay.** Activation curve generated by S2765 (FXa specific chromogenic substrate) as FIX activation is determined indirectly by the activation of FX (10nM) (n=3). FIX activity with rcM^pro^ (1:1 (mol/mol) E:S ratio) or without protease after 3-h incubation are shown as red and black curves, respectively. As a positive control, the activity of FIX is monitored after treatment with 20μM PCPS (50:50) (an activator of FIX) (⁃, n=3). (**E**) **Activation of FXI monitored by enzymatic assay.** Activation curve generated by S2366 (FXIa specific chromogenic substrate) (n=3). FXI activity with rcM^pro^ (1:1 mol/mol E:S ratio) or without protease after 3-h incubation are shown as red and black curves, respectively. As a positive control, the activity of FXIa is monitored (⁃, n=1). (**F**) **Effect of Mpro on ATIII-mediated inhibition of thrombin.** Activation curve generated by S2338 (α-thrombin specific chromogenic substrate) (n=2). α-Thrombin residual activity after 3-h incubation with ATIII, preincubated with (⁃) or without (⁃) rcM^pro^ (1:1 molar ratio) for 1h at 37°C. The activity of α-thrombin without ATIII is monitored as a positive control (⁃, n=2).

**Figure S4. Screening for M^pro^ ability to activate coagulation factor FVII.** (**A**) **FVII titration curve upon incubation with M^pro^** (different time point from 45 to 180 minutes). Activation curve of FVII is generated by MeSO2-D-CHA-But-Arg-pNA (FVIIa specific chromogenic substrate). FVII activity is monitored in the presence of 100 nM recombinant tissue factor (TF) added just before the kinetic measurements. FVII activity in presence of rcM^pro^ (10:1 (mol/mol) E:S ratio) and in absence of rcM^pro^ is shown with different scale of blue and grey based on the incubation time. (**B**) **FVII titration curve with different concentrations of FVII** (0-10nM). Activation curve of FVII is generated by CH_3_SO_2_-D-Cycloehyl-Ala-But-Arg-pNA (FVIIa specific chromogenic substrate). FVIIa activity is monitored in the presence of 100 nM recombinant tissue factor (TF) added just before starting the kinetic measurements.

**Figure S5. Chacterization of rcM^pro^.** (**A**) SDS-PAGE (Bolt Bis-TRIS 4-12% precast gel) analysis of rcM^pro^ with Coomassie staining (Simply Blue SafeStain, Invitrogen). Lane 1: 0.5 µg, lane 2: 1 µg, M: molecular weight protein standards. (**B**) Intact MS analysis of rcM^pro^. M^pro^ signal is indicated by the arrow.

**Figure S6. Cleavage of synthetic peptide substrates by M^pro^, as deduced by RP-HPLC analysis.** Bar plots representing the intensity, of the full-length peptides FTRLQSLNE and FTRLRSLEN and of the corresponding N-terminal and C-terminal fragments generated after cleavage at **Q**↓**S** or **R**↓**S** bonds, at time 0 and after 16-h treatment with (red bars) or without (mock condition, violet bars) rcM^pro^. For each peptide species, the intensity values are expressed as the percent area under the curve of the corresponding chromatographic peak, compared to the total peak area in the chromatogram.

**Figure S7. HDX-MS data of deuterium uptake by rcM^pro^ before and after incubation with PPACK.** (**A**) Coverage map where all the peptides considered for the HDX-MS differential analysis are represented by blue bars under the M^pro^ sequence. (**B**) Quantification of Cys145-underivatized peptides as a function of time in HDX-MS measurements. Peptides 141-150 and 142-150, deriving from the cleavage of M^pro^ by pepsin during HDX-MS analysis, are quantified at different time points (0.25, 1, 5, 30, 120min) in the presence or absence of PPACK. The experiment was performed in triplicate. Each dot represents the results obtained for a biological replicate, with the box plot boundaries indicating the quantiles Q1 (25%) and Q3 (75%) and the median value denoted by a line across the box.

**Figure S8. Targeted mass spectrometry analysis of FVII activation peptides. (A,B)** Analysis of iRT peptides used as an internal standard for the targeted analysis. Retention time and the integration of MS2 fragments are reported in panel A and B. (**C**) Elution profile of FVII activation peptide (Dimethyl)IVGGKVCPK(Dimethyl)GECPWQVLLLVNG. The top 8 most intense MS2 fragments are reported.

## Notes

### Competing Interest Statement

The authors have declared no competing interest.

## References

1. Beristain-Covarrubias, N. et al. Understanding Infection-Induced Thrombosis: Lessons Learned From Animal Models. Frontiers in Immunology 10, (2019).

2. Linder, M., Müller-Berghaus, G., Lasch, H. G. & Gagel, C. Virus infection and blood coagulation. Thrombosis et diathesis haemorrhagica 23, 1–11 (1970).

3. Antoniak, S. & Mackman, N. Multiple roles of the coagulation protease cascade during virus infection. Blood 123, 2605–2613 (2014).

4. Goeijenbier, M. et al. Review: Viral infections and mechanisms of thrombosis and bleeding. Journal of Medical Virology 84, 1680–1696 (2012).

5. Galli, L., Gerdes, V. E. A., Guasti, L. & Squizzato, A. Thrombosis Associated with Viral Hepatitis. J Clin Transl Hepatol 2, 234–239 (2014).

6. Chuang, Y.-C. et al. Factors contributing to the disturbance of coagulation and fibrinolysis in dengue virus infection. J Formos Med Assoc 112, 12–17 (2013).

7. Justo, D., Finn, T., Atzmony, L., Guy, N. & Steinvil, A. Thrombosis associated with acute cytomegalovirus infection: a meta-analysis. Eur J Intern Med 22, 195–199 (2011).

8. Nicholson, A. C. & Hajjar, D. P. Herpesvirus in atherosclerosis and thrombosis: etiologic agents or ubiquitous bystanders? Arterioscler Thromb Vasc Biol 18, 339–348 (1998).

9. Geisbert, T. W. et al. Mechanisms Underlying Coagulation Abnormalities in Ebola Hemorrhagic Fever: Overexpression of Tissue Factor in Primate Monocytes/Macrophages Is a Key Event. The Journal of Infectious Diseases 188, 1618–1629 (2003).

10. Huerta-Zepeda, A. et al. Crosstalk between coagulation and inflammation during Dengue virus infection. Thromb Haemost 99, 936–943 (2008).

11. Turnic, T. N. et al. Bradykinin and Galectin-3 in Survived and Deceased Patients with COVID-19 Pneumonia: An Increasingly Promising Biochemical Target. Oxid Med Cell Longev 2022, 7920915 (2022).

12. Giannis, D., Douketis, J. D. & Spyropoulos, A. C. Anticoagulant therapy for COVID-19: What we have learned and what are the unanswered questions? European Journal of Internal Medicine 96, 13–16 (2022).

13. Gorog, D. A. et al. Current and novel biomarkers of thrombotic risk in COVID-19: a Consensus Statement from the International COVID-19 Thrombosis Biomarkers Colloquium. Nature Reviews Cardiology 2022 19:7 19, 475–495 (2022).

14. Zheng, Y.-Y., Ma, Y.-T., Zhang, J.-Y. & Xie, X. COVID-19 and the cardiovascular system. Nature Reviews Cardiology 2020 17:5 17, 259–260 (2020).

15. Lodigiani, C. et al. Venous and arterial thromboembolic complications in COVID-19 patients admitted to an academic hospital in Milan, Italy. Thrombosis research 191, 9–14 (2020).

16. Helms, J. et al. High risk of thrombosis in patients with severe SARS-CoV-2 infection: a multicenter prospective cohort study. Intensive care medicine 46, 1089–1098 (2020).

17. Llitjos, J. F. et al. High incidence of venous thromboembolic events in anticoagulated severe COVID-19 patients. Journal of thrombosis and haemostasis J: JTH 18, 1743–1746 (2020).

18. Lucas, C. et al. Longitudinal analyses reveal immunological misfiring in severe COVID-19. Nature 2020 584:7821 584, 463–469 (2020).

19. Iwasaki, M. et al. Inflammation Triggered by SARS-CoV-2 and ACE2 Augment Drives Multiple Organ Failure of Severe COVID-19: Molecular Mechanisms and Implications. Inflammation 2020 44:1 44, 13–34 (2020).

20. Mahmudpour, M., Roozbeh, J., Keshavarz, M., Farrokhi, S. & Nabipour, I. COVID-19 cytokine storm: The anger of inflammation. Cytokine 133, (2020).

21. Hu, B., Huang, S. & Yin, L. The cytokine storm and COVID-19. Journal of medical virology 93, 250–256 (2021).

22. Wolf, A. et al. The mechanistic basis linking cytokine storm to thrombosis in COVID-19. Thrombosis Update 8, 100110 (2022).

23. Esmon, C. T. Possible involvement of cytokines in diffuse intravascular coagulation and thrombosis. Baillieres Best Pract Res Clin Haematol 12, 343–359 (1999).

24. Rex, D. A. B., Vaid, N., Deepak, K., Dagamajalu, S. & Prasad, T. S. K. A comprehensive review on current understanding of bradykinin in COVID-19 and inflammatory diseases. Molecular biology reports 49, (2022).

25. Stojanovski, B. M. & Di Cera, E. Comparative sequence analysis of vitamin K-dependent coagulation factors. Journal of Thrombosis and Haemostasis 20, 2837–2849 (2022).

26. Hoffman, M. Remodeling the blood coagulation cascade. Journal of thrombosis and thrombolysis 16, 17–20 (2003).

27. Pontarollo, G. et al. Non-canonical proteolytic activation of human prothrombin by subtilisin from Bacillus subtilis may shift the procoagulant-anticoagulant equilibrium toward thrombosis. The Journal of biological chemistry 292, 15161–15179 (2017).

28. Malone, B., Urakova, N., Snijder, E. J. & Campbell, E. A. Structures and functions of coronavirus replication–transcription complexes and their relevance for SARS-CoV-2 drug design. Nat Rev Mol Cell Biol 23, 21–39 (2022).

29. Moustaqil, M. et al. SARS-CoV-2 proteases PLpro and 3CLpro cleave IRF3 and critical modulators of inflammatory pathways (NLRP12 and TAB1): implications for disease presentation across species. Emerging Microbes and Infections 10, 178–195 (2021).

30. Pablos, I. et al. Mechanistic insights into COVID-19 by global analysis of the SARS-CoV-2 3CLpro substrate degradome. Cell Reports 37, 109892 (2021).

31. Meyer, B. et al. Characterising proteolysis during SARS-CoV-2 infection identifies viral cleavage sites and cellular targets with therapeutic potential. Nature Communications 2021 12:1 12, 1–16 (2021).

32. Wenzel, J. et al. The SARS-CoV-2 main protease Mpro causes microvascular brain pathology by cleaving NEMO in brain endothelial cells. Nat Neurosci 24, 1522–1533 (2021).

33. Martínez-Fleta, P. et al. SARS-CoV-2 Cysteine-like Protease Antibodies Can Be Detected in Serum and Saliva of COVID-19-Seropositive Individuals. Journal of immunology (Baltimore, Md. J: 1950) 205, 3130–3140 (2020).

34. Jiang, H. wei et al. SARS-CoV-2 proteome microarray for global profiling of COVID-19 specific IgG and IgM responses. Nature communications 11, (2020).

35. Wang, W. et al. Detection of SARS-CoV-2 in Different Types of Clinical Specimens. JAMA 323, 1843–1844 (2020).

36. Gordon, D. E. et al. A SARS-CoV-2 protein interaction map reveals targets for drug repurposing. Nature 583, 459–468 (2020).

37. Zhou, Y. et al. A comprehensive SARS-CoV-2–human protein–protein interactome reveals COVID-19 pathobiology and potential host therapeutic targets. Nature Biotechnology 2022 41:1 41, 128–139 (2022).

38. Liu, X. et al. SARS-CoV-2–host proteome interactions for antiviral drug discovery. Molecular Systems Biology 17, e10396 (2021).

39. Biembengut, Í. V. & de Souza, T. de A. C. B. Coagulation modifiers targeting SARS-CoV-2 main protease Mpro for COVID-19 treatment: an in silico approach. Memorias do Instituto Oswaldo Cruz 115, 1–4 (2020).

40. Zhang, L. et al. Crystal structure of SARS-CoV-2 main protease provides a basis for design of improved α-ketoamide inhibitors. Science (New York, N.Y.) 368, 409–412 (2020).

41. Fang, S., Shen, H., Wang, J., Tay, F. P. L. & Liu, D. X. Functional and Genetic Studies of the Substrate Specificity of Coronavirus Infectious Bronchitis Virus 3C-Like Proteinase. Journal of Virology 84, 7325–7336 (2010).

42. Shaqra, A. M. et al. Defining the substrate envelope of SARS-CoV-2 main protease to predict and avoid drug resistance. Nature Communications 2022 13:1 13, 1–11 (2022).

43. Zhao, Y. et al. Structural basis for replicase polyprotein cleavage and substrate specificity of main protease from SARS-CoV-2. Proceedings of the National Academy of Sciences of the United States of America 119, e2117142119 (2022).

44. Rut, W. et al. SARS-CoV-2 Mpro inhibitors and activity-based probes for patient-sample imaging. Nature Chemical Biology 2020 17:2 17, 222–228 (2020).

45. Koudelka, T., et al. N-Terminomics for the Identification of In Vitro Substrates and Cleavage Site Specificity of the SARS-CoV-2 Main Protease. Proteomics 21, (2021).

46. Dai, W. et al. Structure-based design of antiviral drug candidates targeting the SARS-CoV-2 main protease. Science 368, 1331–1335 (2020).

47. Owen, D. R. et al. An oral SARS-CoV-2 Mpro inhibitor clinical candidate for the treatment of COVID-19. Science 374, 1586–1593 (2021).

48. Fornasier, E. et al. A new inactive conformation of SARS-CoV-2 main protease. Acta crystallographica. Section D, Structural biology 78, 363–378 (2022).

49. Adcock DM, Hoefner DM, Kottke-Marchant K, Marlar RA, Szamosi DI, W. DJ. H21-A5 Collection, Transport, and Processing of Blood Specimens for Testing Plasma-Based Coagulation Assays and Molecular Hemostasis Assays; Approved Guideline-Fifth Edition. (2008).

50. Mui, B., Chow, L. & Hope, M. J. Extrusion Technique to Generate Liposomes of Defined Size. Methods in Enzymology 367, 3–14 (2003).

51. Bouck, E. G., Zunica, E. R. & Nieman, M. T. Optimizing the presentation of bleeding and thrombosis data:Responding to censored data using Kaplan-Meier curves. Thrombosis research 158, 154 (2017).

52. Therneau, T. A package for survival analysis in R.

53. Kassambara, A., Kosinski, M. & Biecek, P. survminer: Drawing Survival Curves using ‘ggplot2’. 0.4.9 10.32614/CRAN.package.survminer (2016).

54. Uliana, F. et al. Mapping specificity, cleavage entropy, allosteric changes and substrates of blood proteases in a high-throughput screen. Nature communications 12, (2021).

55. Yan, B. et al. Analysis of post-translational modifications in recombinant monoclonal antibody IgG1 by reversed-phase liquid chromatography/mass spectrometry. Journal of chromatography. A 1164, 153–161 (2007).

56. Wiśniewski, J. R. Filter-Aided Sample Preparation for Proteome Analysis. Methods Mol Biol 1841, 3–10 (2018).

57. Acquasaliente, L., Pierangelini, A., Pagotto, A., Pozzi, N. & De Filippis, V. From Haemadin to Haemanorm: Synthesis and Characterization of Full-length Haemadin from the leech Haemadipsa sylvestris and of a Novel Bivalent, Highly Potent Thrombin Inhibitor (Haemanorm). Protein Sci e4825 (2023) doi:10.1002/pro.4825.

58. Kuzmič, P. History, variants and usage of the ‘Morrison equation’ in enzyme inhibition kinetics.

59. Peterle, D. et al. A serine protease secreted from Bacillus subtilis cleaves human plasma transthyretin to generate an amyloidogenic fragment. Commun Biol 3, 764 (2020).

60. Kleifeld, O. et al. Identifying and quantifying proteolytic events and the natural N terminome by terminal amine isotopic labeling of substrates. Nature protocols 6, 1578–1611 (2011).

61. Rappsilber, J., Mann, M. & Ishihama, Y. Protocol for micro-purification, enrichment, pre-fractionation and storage of peptides for proteomics using StageTips. Nat Protoc 2, 1896– 1906 (2007).

62. MacLean, B., et al. Skyline: an open source document editor for creating and analyzing targeted proteomics experiments. Bioinformatics 26, 966–968 (2010).

63. Weisel, J. W. & Nagaswami, C. Computer modeling of fibrin polymerization kinetics correlated with electron microscope and turbidity observations: clot structure and assembly are kinetically controlled. Biophysical Journal 63, 111–128 (1992).

64. De Cristofaro, R. & Di Cera, E. Phenomenological analysis of the clotting curve. Journal of Protein Chemistry 1991 10:5 10, 455–468 (1991).

65. Kattula, S., Byrnes, J. R. & Wolberg, A. S. Fibrinogen and Fibrin in Hemostasis and Thrombosis. Arteriosclerosis, Thrombosis, and Vascular Biology 37, e13–e21 (2017).

66. Pieters, M., Jerling, J. C. & Weisel, J. W. Effect of freeze-drying, freezing and frozen storage of blood plasma on fibrin network characteristics. Thrombosis Research 107, 263–269 (2002).

67. Farkas, Á. Z. et al. Structure, Mechanical, and Lytic Stability of Fibrin and Plasma Coagulum Generated by Staphylocoagulase From Staphylococcus aureus. Front Immunol 10, 2967 (2019).

68. Bernardi, F. & Mariani, G. Biochemical, molecular and clinical aspects of coagulation factor VII and its role in hemostasis and thrombosis. Haematologica 106, 351–362 (2021).

69. Björk, I. & Lindahl, U. Mechanism of the anticoagulant action of heparin. Mol Cell Biochem 48, 161–182 (1982).

70. Mitropoulos, K., Martin, J., Reeves, B. & Esnouf, M. The activation of the contact phase of coagulation by physiologic surfaces in plasma: the effect of large negatively charged liposomal vesicles. Blood 73, 1525–1533 (1989).

71. KE, K. & Morrissey, J. The activation of factor VII by a variety of potential activators. The FASEB Journal 27, 995.6-995.6 (2013).

72. Naudin, C., Burillo, E., Blankenberg, S., Butler, L. & Renné, T. Factor XII Contact Activation. Seminars in thrombosis and hemostasis 43, 814–826 (2017).

73. Kenne, E. & Renné, T. Factor XII: A drug target for safe interference with thrombosis and inflammation. Drug Discovery Today 19, 1459–1464 (2014).

74. Chuck, C.-P., Chow, H.-F., Wan, D. C.-C. & Wong, K.-B. Profiling of Substrate Specificities of 3C-Like Proteases from Group 1, 2a, 2b, and 3 Coronaviruses. PLOS ONE 6, e27228 (2011).

75. Hegyi, A. & Ziebuhr, J. Conservation of substrate specificities among coronavirus main proteases. The Journal of general virology 83, 595–599 (2002).

76. Schechter, I. & Berger, A. On the size of the active site in proteases. I. Papain. Biochemical and Biophysical Research Communications 27, 157–162 (1967).

77. Scott, B. M., Lacasse, V., Blom, D. G., Tonner, P. D. & Blom, N. S. Predicted coronavirus Nsp5 protease cleavage sites in the human proteome. BMC Genomic Data 23, 25 (2022).

78. Anand, K. et al. Structure of coronavirus main proteinase reveals combination of a chymotrypsin fold with an extra alpha-helical domain. EMBO J 21, 3213–3224 (2002).

79. Kovach, I. M., Kelley, P., Eddy, C., Jordan, F. & Baykal, A. Proton Bridging in the Interactions of Thrombin with Small Inhibitors. Biochemistry 48, 7296–7304 (2009).

80. Engen, J. R. & Komives, E. A. Complementarity of Hydrogen/Deuterium Exchange Mass Spectrometry and Cryo-Electron Microscopy. Trends in Biochemical Sciences 45, 906–918 (2020).

81. Masson, G. R. et al. Recommendations for performing, interpreting and reporting hydrogen deuterium exchange mass spectrometry (HDX-MS) experiments. Nat Methods 16, 595–602 (2019).

82. Konermann, L., Vahidi, S. & Sowole, M. A. Mass Spectrometry Methods for Studying Structure and Dynamics of Biological Macromolecules. Anal. Chem. 86, 213–232 (2014).

83. Zhang, L. et al. Crystal structure of SARS-CoV-2 main protease provides a basis for design of improved α-ketoamide inhibitors. Science 368, 409–412 (2020).

84. Domon, B. & Aebersold, R. Options and considerations when selecting a quantitative proteomics strategy. Nat Biotechnol 28, 710–721 (2010).

85. Augustsson, C. & Persson, E. In vitro evidence of a tissue factor-independent mode of action of recombinant factor VIIa in hemophilia. Blood 124, 3172–3174 (2014).

86. Einarson, T. R., Acs, A., Ludwig, C. & Panton, U. H. Prevalence of cardiovascular disease in type 2 diabetes: a systematic literature review of scientific evidence from across the world in 2007-2017. Cardiovasc Diabetol 17, 83 (2018).

87. Luke, R. G. Chronic renal failure--a vasculopathic state. N Engl J Med 339, 841–843 (1998).

88. Lin, H. et al. Epidemiology and Risk Factors of Portal Venous System Thrombosis in Patients With Inflammatory Bowel Disease: A Systematic Review and Meta-Analysis. Front Med (Lausanne) 8, 744505 (2021).

89. Patmore, S., Dhami, S. P. S. & O’Sullivan, J. M. Von Willebrand factor and cancer; metastasis and coagulopathies. J Thromb Haemost 18, 2444–2456 (2020).

90. Sokolov, A. V. et al. Thrombin inhibits the anti-myeloperoxidase and ferroxidase functions of ceruloplasmin: relevance in rheumatoid arthritis. Free Radic Biol Med 86, 279–294 (2015).

91. Pozzi, N. & Acquasaliente, L. β2 –Glycoprotein I binds to thrombin and selectively inhibits the enzyme procoagulant functions – PubMed. https://pubmed.ncbi.nlm.nih.gov/23578283/.

92. Meroni, P. L., Borghi, M. O., Raschi, E. & Tedesco, F. Pathogenesis of antiphospholipid syndrome: understanding the antibodies. Nat Rev Rheumatol 7, 330–339 (2011).

93. Acquasaliente, L. Molecular mapping of α-thrombin (αT)/β2-glycoprotein I (β2GpI) interaction reveals how β2GpI affects αT functions – PubMed. https://pubmed.ncbi.nlm.nih.gov/27760842/.

94. Zamolodchikov, D., Renné, T. & Strickland, S. The Alzheimer’s disease peptide β-amyloid promotes thrombin generation through activation of coagulation factor XII. J Thromb Haemost 14, 995–1007 (2016).

95. Bever, K. M. et al. Risk factors for venous thromboembolism in immunoglobulin light chain amyloidosis. Haematologica 101, 86–90 (2016).

96. Ruberg, F. L., Grogan, M., Hanna, M., Kelly, J. W. & Maurer, M. S. Transthyretin Amyloid Cardiomyopathy: JACC State-of-the-Art Review. J Am Coll Cardiol 73, 2872–2891 (2019).

97. Acquasaliente, L. & De Filippis, V. The Role of Proteolysis in Amyloidosis. Int J Mol Sci 24, 699 (2022).

98. Acquasaliente, L. et al. Exogenous human α-Synuclein acts in vitro as a mild platelet antiaggregant inhibiting α-thrombin-induced platelet activation. Sci Rep 12, 9880 (2022).

99. Levi, M. & Ten Cate, H. Disseminated intravascular coagulation. N Engl J Med 341, 586– 592 (1999).

100. Loof, T. G. et al. Staphylococcus aureus-induced clotting of plasma is an immune evasion mechanism for persistence within the fibrin network. Microbiology (Reading) 161, 621–627 (2015).

101. Iba, T., Levy, J. H., Levi, M. & Thachil, J. Coagulopathy in COVID-19. J Thromb Haemost 18, 2103–2109 (2020).

102. Lancellotti, S. et al. Formation of methionine sulfoxide by peroxynitrite at position 1606 of von Willebrand factor inhibits its cleavage by ADAMTS-13: A new prothrombotic mechanism in diseases associated with oxidative stress. Free Radic Biol Med 48, 446–456 (2010).

103. Lancellotti, S. et al. Oxidized von Willebrand factor is efficiently cleaved by serine proteases from primary granules of leukocytes: divergence from ADAMTS-13. J Thromb Haemost 9, 1620–1627 (2011).

104. De Filippis, V., et al. Oxidation of Met1606 in von Willebrand factor is a risk factor for thrombotic and septic complications in chronic renal failure. Biochem J 442, 423–432 (2012).

105. Pontarollo, G. et al. Non-canonical proteolytic activation of human prothrombin by subtilisin from Bacillus subtilis may shift the procoagulant-anticoagulant equilibrium toward thrombosis. J Biol Chem 292, 15161–15179 (2017).

106. Bikdeli, B. et al. COVID-19 and Thrombotic or Thromboembolic Disease: Implications for Prevention, Antithrombotic Therapy, and Follow-Up: JACC State-of-the-Art Review. Journal of the American College of Cardiology 75, 2950–2973 (2020).

107. Conway, E. M. et al. Understanding COVID-19-associated coagulopathy. Nat Rev Immunol 22, 639–649 (2022).

108. Hofman, Z., de Maat, S., Hack, C. E. & Maas, C. Bradykinin: Inflammatory Product of the Coagulation System. Clin Rev Allergy Immunol 51, 152–161 (2016).

109. Reynolds, N. D. et al. The SARS-CoV-2 SSHHPS Recognized by the Papain-like Protease. ACS Infect Dis 7, 1483–1502 (2021).

110. Andersson, M. I. et al. SARS-CoV-2 RNA detected in blood products from patients with COVID-19 is not associated with infectious virus. Wellcome Open Res 5, 181 (2020).

111. Shamanaev, A. et al. Model for surface-dependent factor XII activation: the roles of factor XII heavy chain domains. Blood Adv 6, 3142–3154 (2022).

112. Versteeg, H. H., Heemskerk, J. W. M., Levi, M. & Reitsma, P. H. New Fundamentals in Hemostasis. Physiological Reviews 93, 327–358 (2013).

113. Yong, J. & Toh, C.-H. The convergent model of coagulation. Journal of Thrombosis and Haemostasis 22, 2140–2146 (2024).

114. Owen, M. J. et al. Mathematical models of coagulation—are we there yet? Journal of Thrombosis and Haemostasis 22, 1689–1703 (2024).

115. Ziliotto, N. et al. Coagulation Factor XII Levels and Intrinsic Thrombin Generation in Multiple Sclerosis. Front Neurol 9, 245 (2018).

