## Supplementary file for "The Main protease (M^pro^) from SARS-CoV-2 triggers plasma clotting *in vitro* by activating coagulation factors VII and FXII"

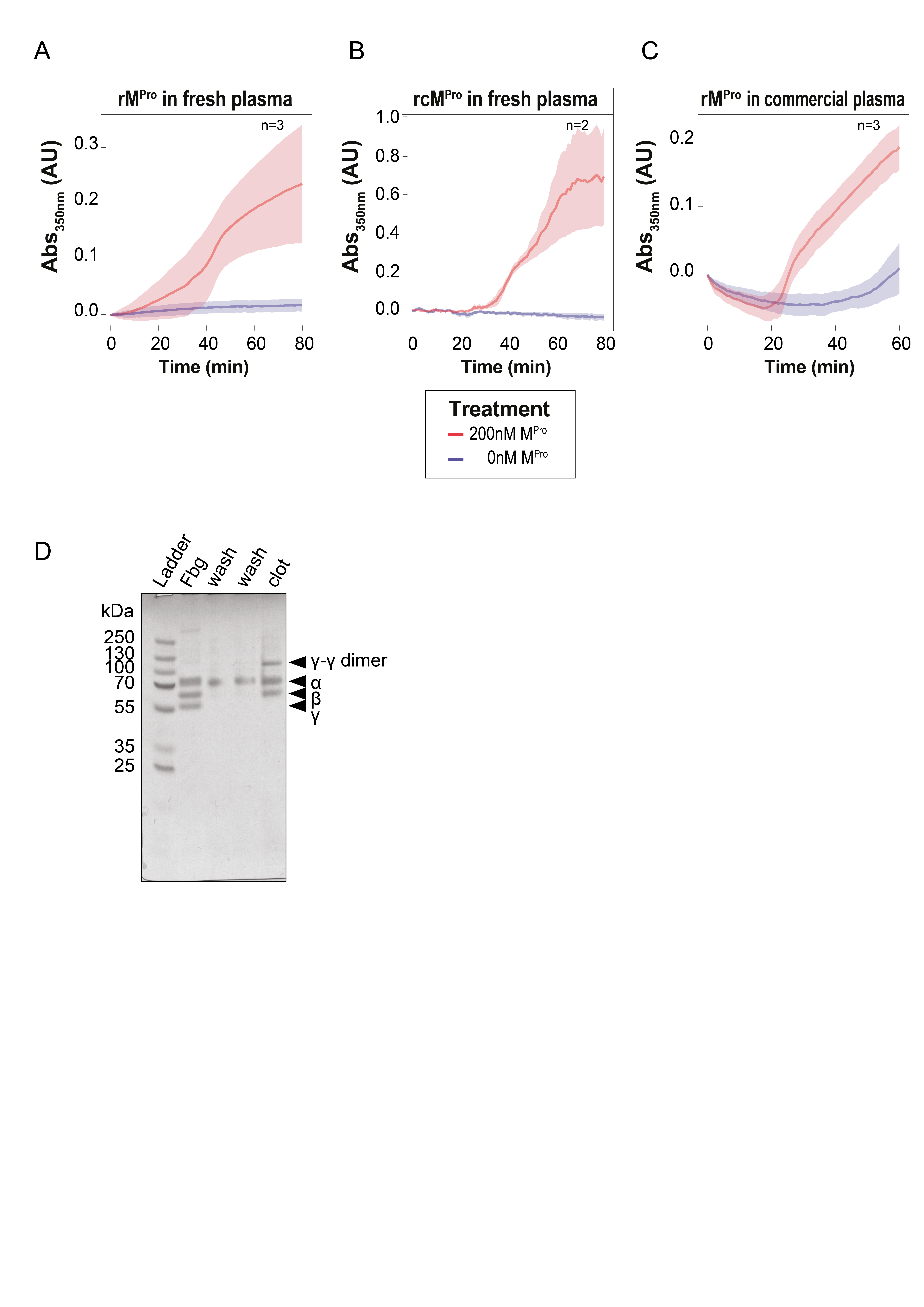

**Figure S1**. **Effect of M^pro^ on fibrin generation in human plasma.**

(**A-C**) Clotting curves obtained after adding rM^pro^ (**A,C**) or rcM^pro^ (**B**) to human plasma samples collected from healthy donors (**A,B**) or reconstituted commercial plasma (**C**). Plasma samples were diluted 1:2 with HBS, re-calcified, and the increase of turbidity at 350 nm was measured over time at 37±0.1°C. Measurements were carried out in triplicate on a single plasma sample with shaded areas corresponding to the standard deviation ±SD at each time point. (**D**) Electrophoretic analysis of plasma clotting reaction. After 1-h incubation at 37°C of 1:2 diluted plasma with rM^pro^ (200 nM), the clot was collected, washed with HBS buffer and resuspended with Gdn-HCl 8M (incubated for 1h at 37°C). Albumin was then removed by treatment with EtOH to a final concentration of 42% for 1h at 4°C and following centrifugation at 16,000g for 45min at 4°C. After removing the supernatant, the pellet was resuspended with SDS sample loading buffer and analysed by reducing SDS-PAGE (Bolt Bis-TRIS 4-12% precast gel) and Coomassie staining (Simply Blue SafeStain, Invitrogen). For comparison, commercial fibrinogen (Fbg) was also loaded. The typical α-, β- and γ-chain of Fbg are indicated by arrows, along with the γ-chain dimer.

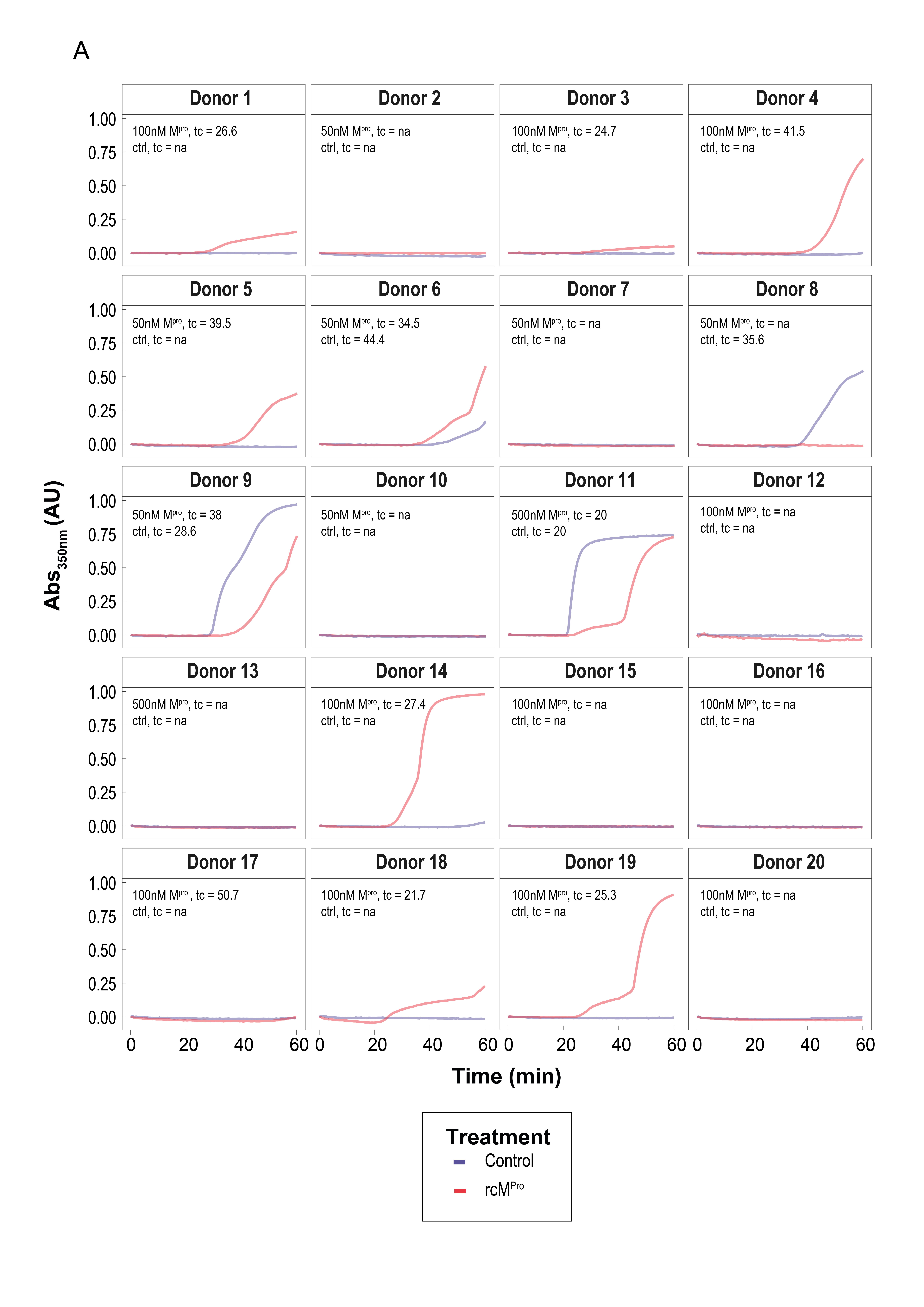

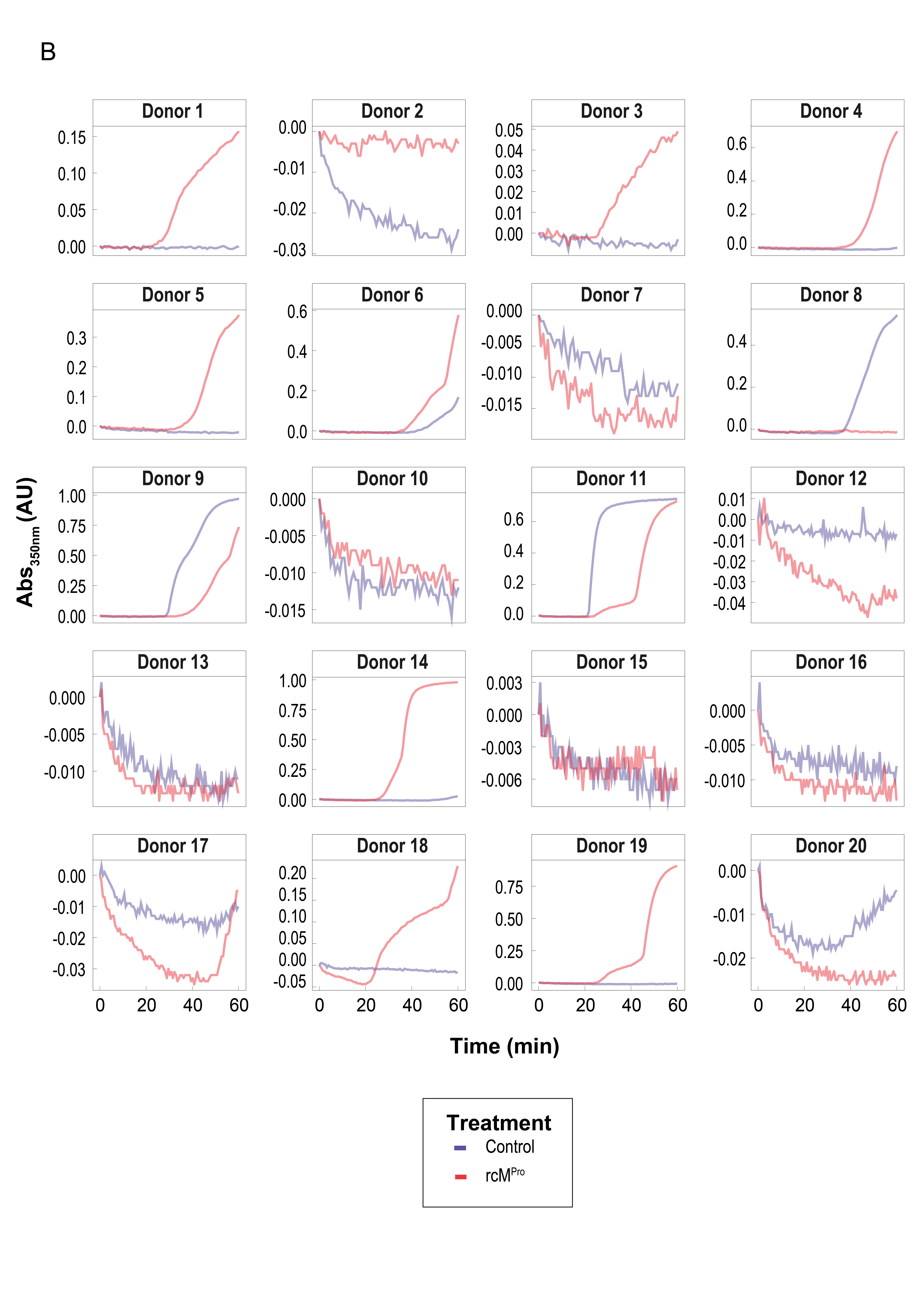

**Figure S2**. **Turbidimetric analysis of the effect of M^pro^ on fibrin generation in human plasma samples from healthy subjects.** Human plasma samples were collected from healthy donors, diluted 1:2 with HBS, and the increase of turbidity at 350 nm was measured over time at 37±0.1°C in the absence and presence of rcM^pro^, at the indicated concentrations. The curves are shown with the same scale (**Figure S2A**) and with a scale normalized for the maximum absorbance changes (**Figure S2B**) to facilitate the identification of sigmoidal profiles.

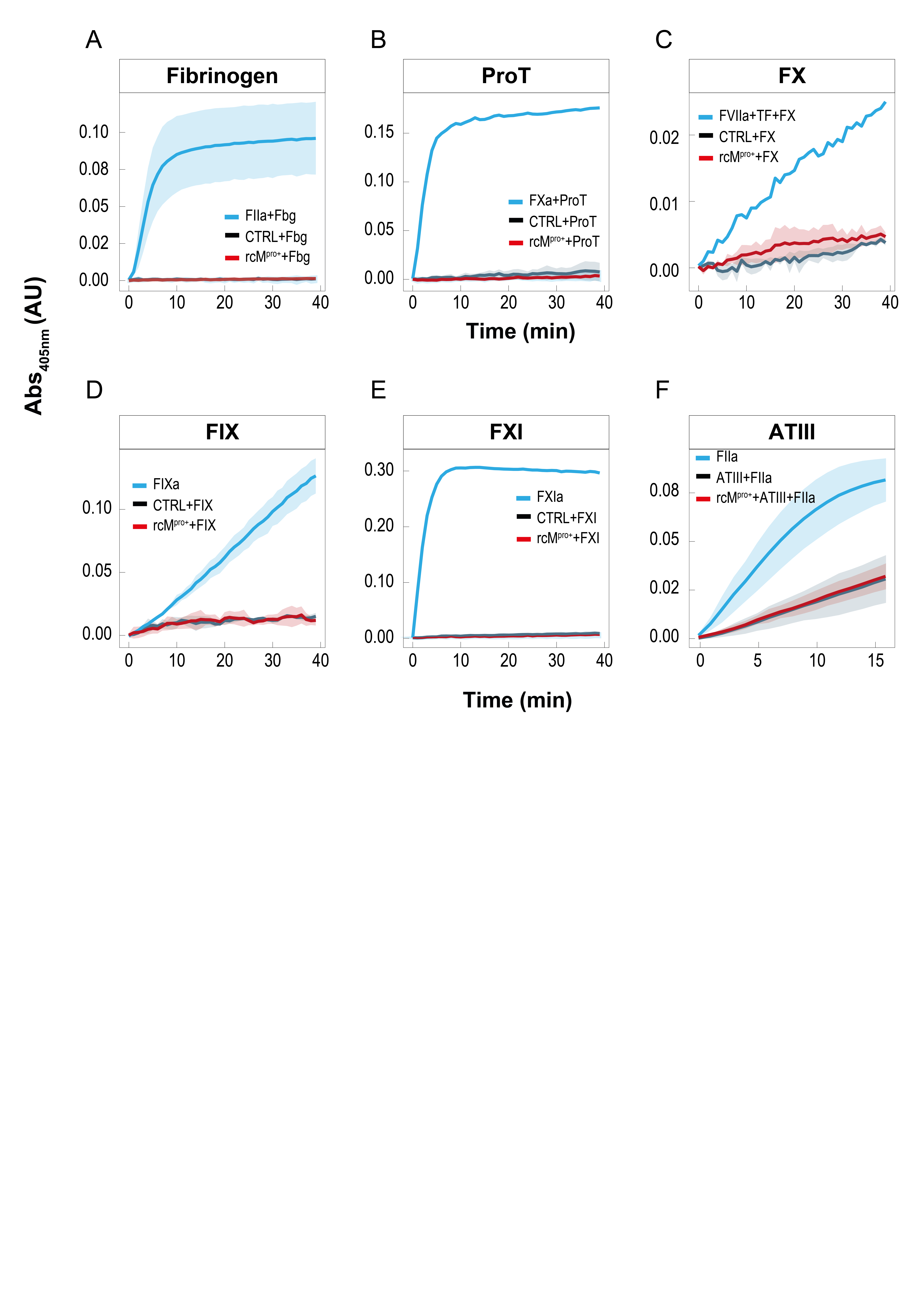

**Figure S3. Screening for M^pro^ ability to activate coagulation factors.**

**(A) Activation of fibrinogen monitored by turbidimetric assay.** Clotting curves (n=2) obtained by adding α-thrombin (2nM) (**▬**) or rcM^pro^ (100 nM) (**▬**) to isolated fibrinogen solution (0.15mg/ml) in the presence of 1.5 mM CaCl_2_. (**B) Activation of prothrombin (FII) monitored by enzymatic assay.** Activation curve generated by S2238 (α-thrombin specific chromogenic substrate) (n=3). Prothrombin activity with rcM^pro^ (10:1 mol/mol E:S ratio) or without protease after 3-h incubation are shown as red and black curves, respectively. As positive control the activity of prothrombin is monitored after treatment with 1nM FXa in the presence of 100μM PCPS (50:50) (**▬**, n=1). (**C**) **Activation of FX monitored by enzymatic assay**. Activation curve generated by S2765 (FXa specific chromogenic substrate) (n=2). FX activity with rcM^pro^ (10:1 (mol/mol) E:S ratio) or without protease after 3-h incubation are shown as red and black curves, respectively. As a positive control, the activity of FX is monitored after treatment with 0.2nM FVIIa in the presence of 2nM recombinant tissue factor (TF) (**▬**, n=1). (**D**) **Activation of FIX monitored by enzymatic assay.** Activation curve generated by S2765 (FXa specific chromogenic substrate) as FIX activation is determined indirectly by the activation of FX (10nM) (n=3). FIX activity with rcM^pro^ (1:1 (mol/mol) E:S ratio) or without protease after 3-h incubation are shown as red and black curves, respectively. As a positive control, the activity of FIX is monitored after treatment with 20μM PCPS (50:50) (an activator of FIX) (**▬**, n=3). (**E**) **Activation of FXI monitored by enzymatic assay.** Activation curve generated by S2366 (FXIa specific chromogenic substrate) (n=3). FXI activity with rcM^pro^ (1:1 mol/mol E:S ratio) or without protease after 3-h incubation are shown as red and black curves, respectively. As a positive control, the activity of FXIa is monitored (**▬**, n=1). (**F**) **Effect of Mpro on ATIII-mediated inhibition of thrombin.** Activation curve generated by S2338 (α-thrombin specific chromogenic substrate) (n=2). α-Thrombin residual activity after 3-h incubation with ATIII, preincubated with (**▬**) or without (**▬**) rcM^pro^ (1:1 molar ratio) for 1h at 37°C. The activity of α-thrombin without ATIII is monitored as a positive control (**▬**, n=2).

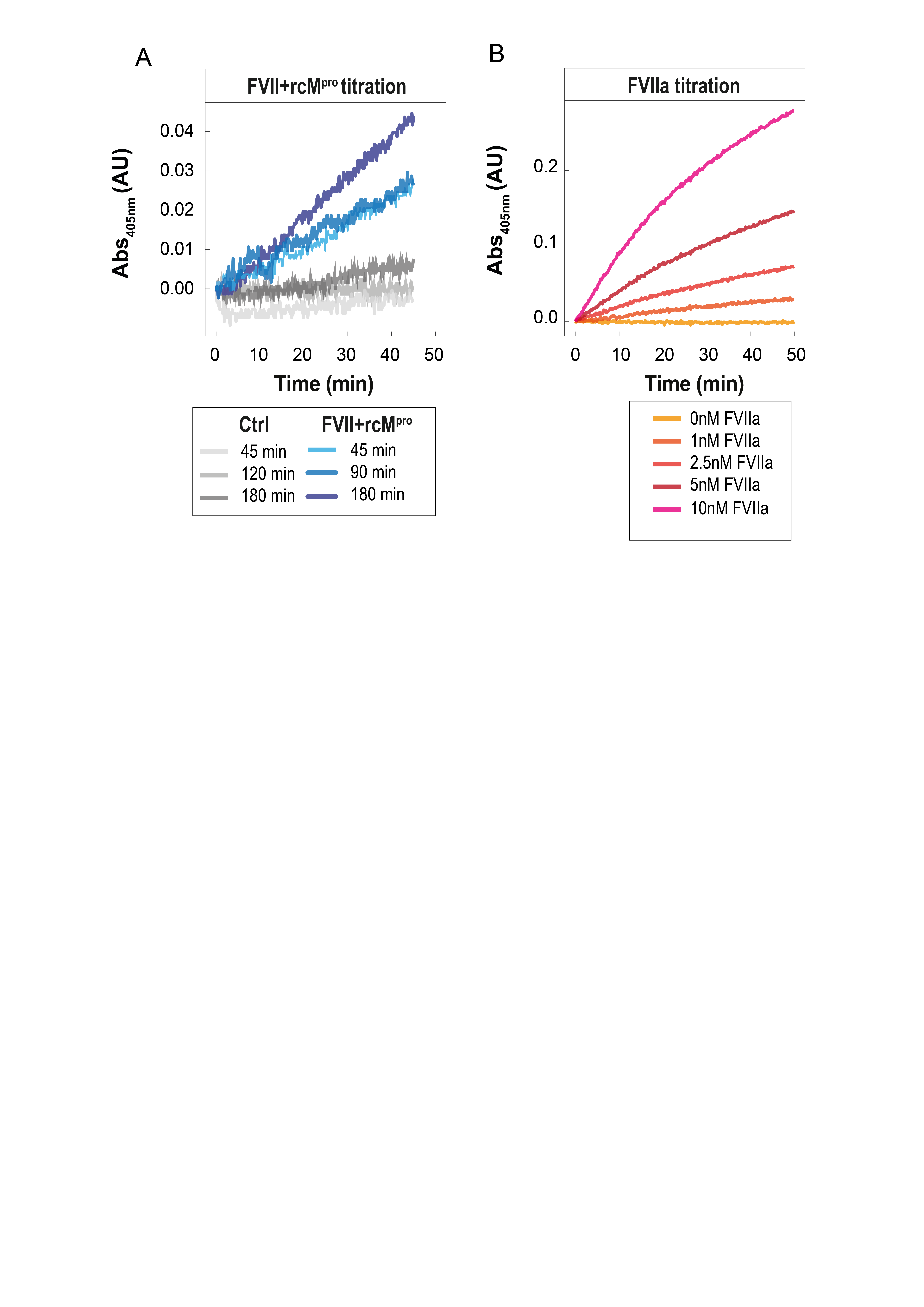

**Figure S4. Screening for M^pro^ ability to activate coagulation factor FVII.**

(**A**) **FVII** **titration curve upon incubation with M^pro^** (different time point from 45 to 180 minutes). Activation curve of FVII is generated by MeSO2-D-CHA-But-Arg-pNA (FVIIa specific chromogenic substrate). FVII activity is monitored in the presence of 100 nM recombinant tissue factor (TF) added just before the kinetic measurements. FVII activity in presence of rcM^pro^ (10:1 (mol/mol) E:S ratio) and in absence of rcM^pro^ is shown with different scale of blue and grey based on the incubation time. (**B**) **FVII** **titration curve with different concentrations of FVII** (0-10nM)**.** Activation curve of FVII is generated by CH_3_SO_2_-D-Cycloehyl-Ala-But-Arg-pNA (FVIIa specific chromogenic substrate). FVIIa activity is monitored in the presence of 100 nM recombinant tissue factor (TF) added just before starting the kinetic measurements.

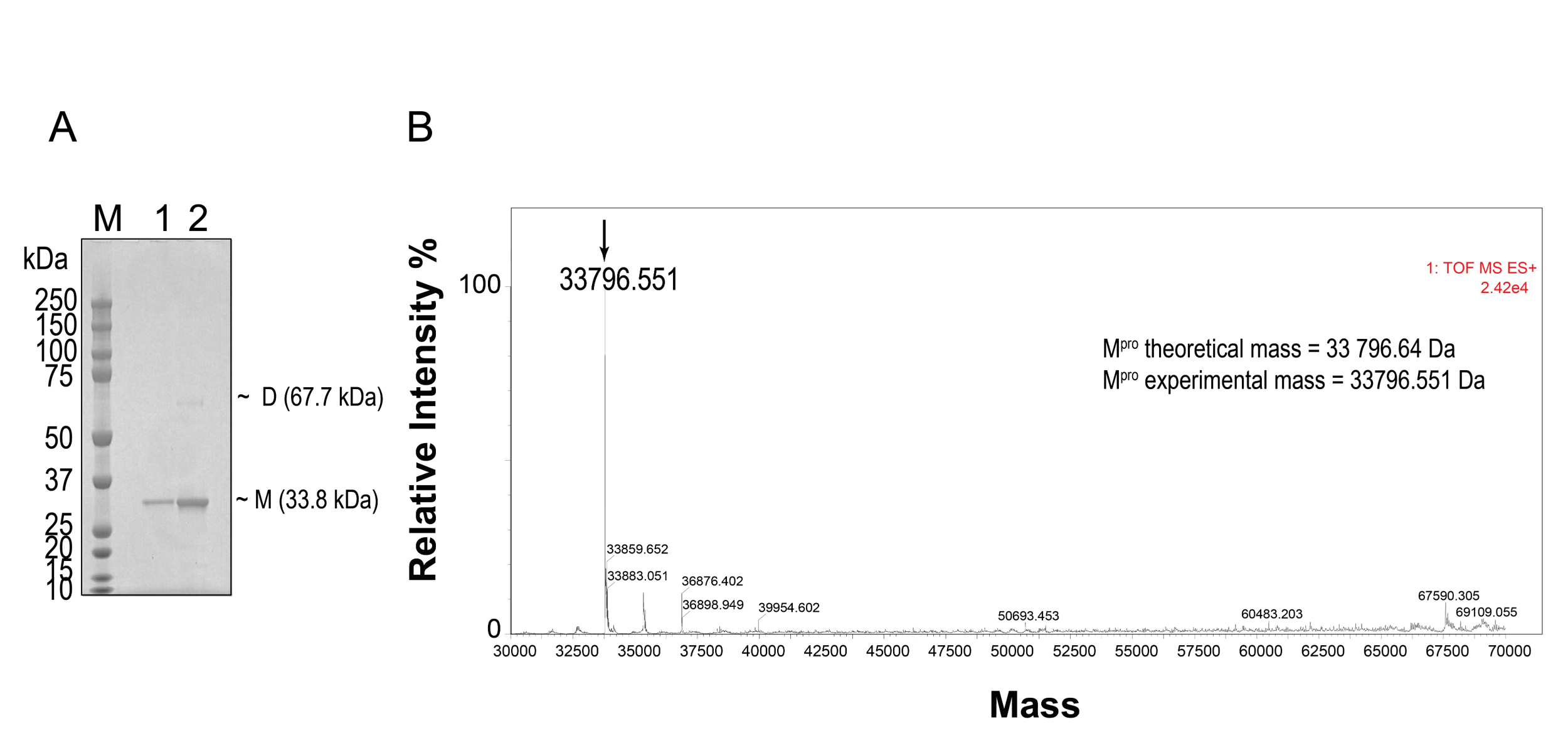

**Figure S5. Chacterization of rcM^pro^.** (**A**) SDS-PAGE (Bolt Bis-TRIS 4-12% precast gel) analysis of rcM^pro^ with Coomassie staining (Simply Blue SafeStain, Invitrogen). Lane 1: 0.5 µg, lane 2: 1 µg, M: molecular weight protein standards. (**B**) Intact MS analysis of rcM^pro^. M^pro^ signal is indicated by the arrow.

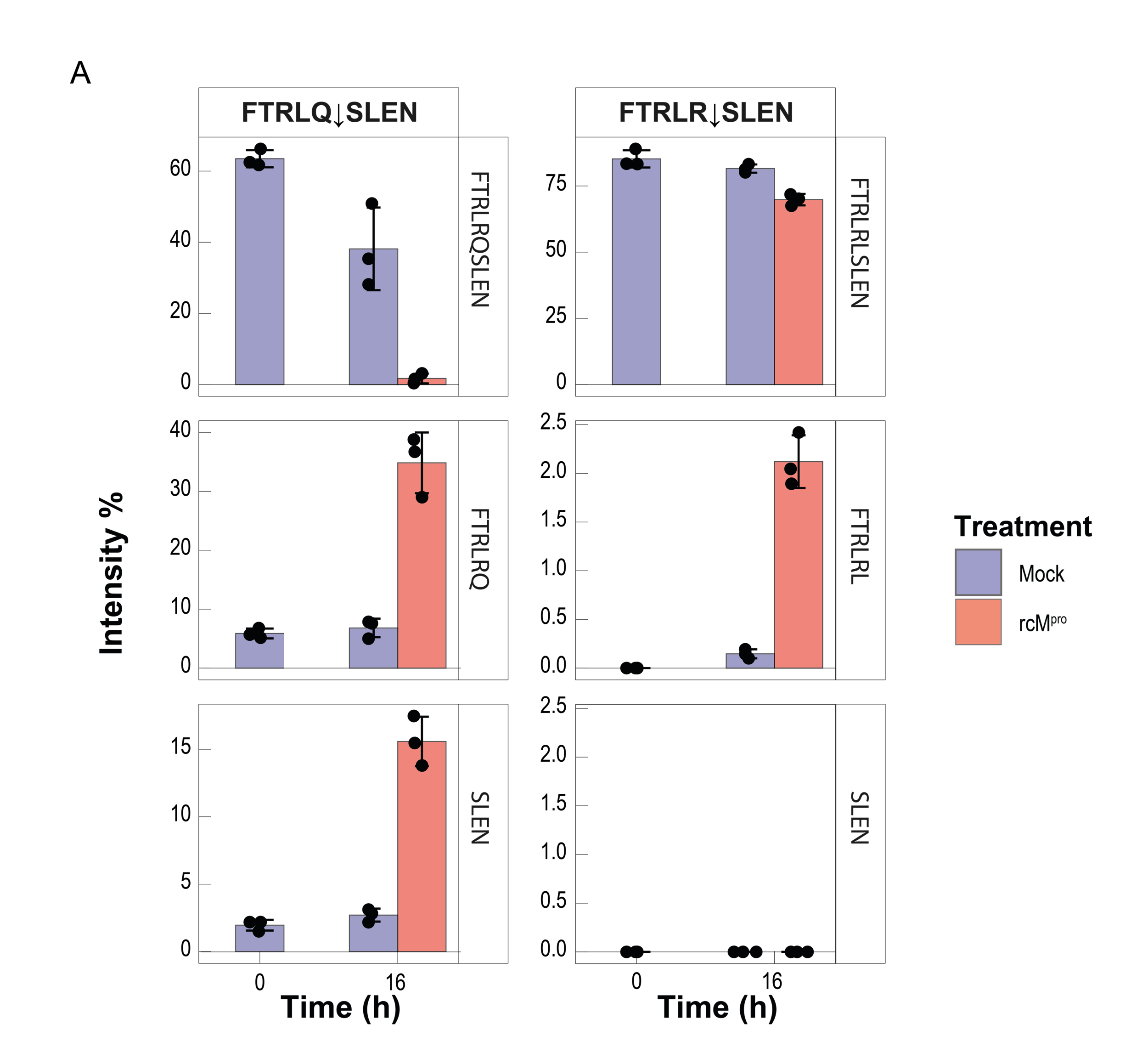

**Figure S6. Cleavage of synthetic peptide substrates by M^pro^, as deduced by RP-HPLC analysis.** Bar plots representing the intensity, of the full-length peptides FTRLQSLNE and FTRLRSLEN and of the corresponding N-terminal and C-terminal fragments generated after cleavage at **Q↓S** or **R↓S** bonds, at time 0 and after 16-h treatment with (red bars) or without (mock condition, violet bars) rcM^pro^. For each peptide species, the intensity values are expressed as the percent area under the curve of the corresponding chromatographic peak, compared to the total peak area in the chromatogram.

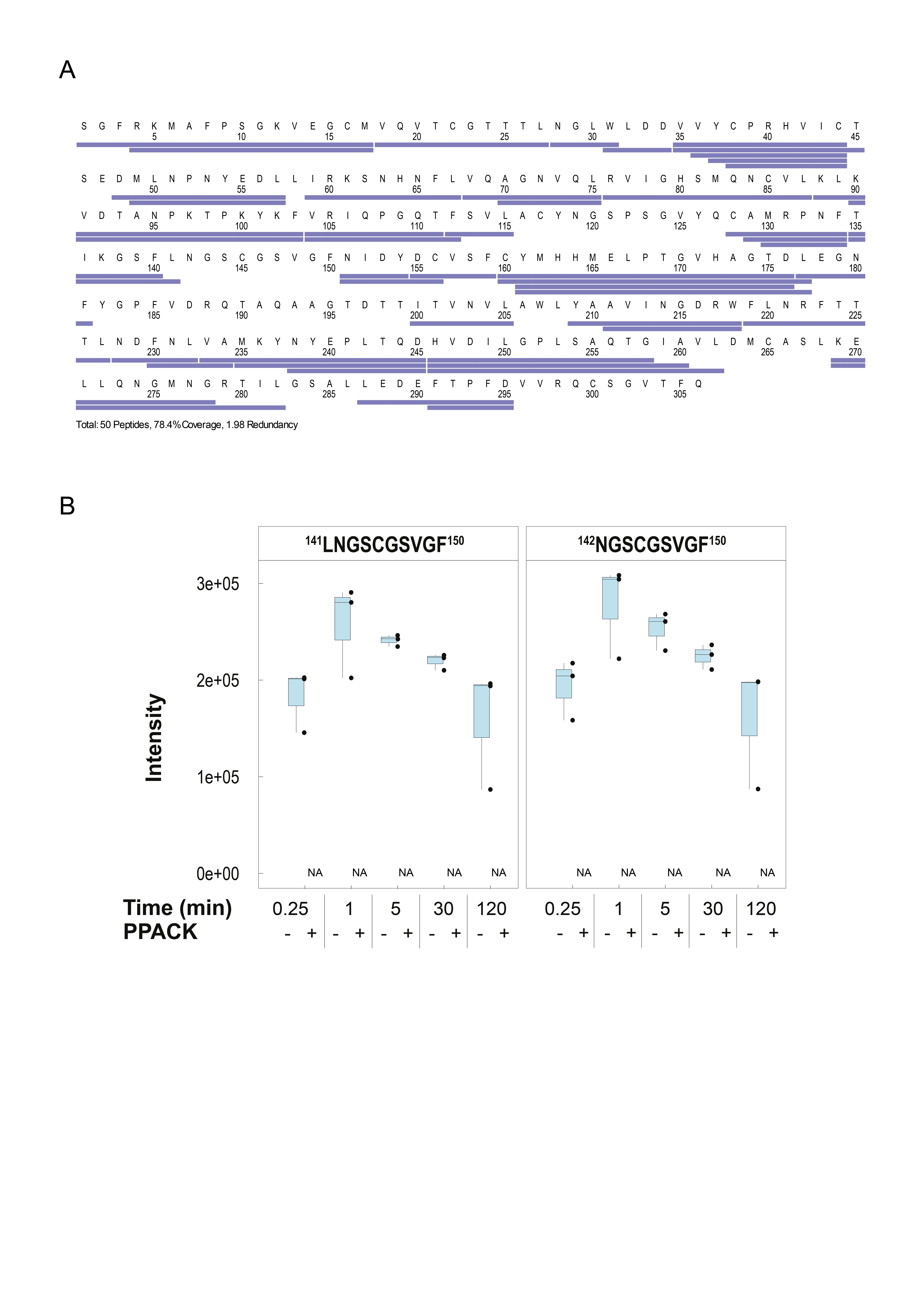

**Figure S7. HDX-MS data of deuterium uptake by rcM^pro^ before and after incubation with PPACK.** (**A**) Coverage map where all the peptides considered for the HDX-MS differential analysis are represented by blue bars under the M^pro^ sequence. (**B**) Quantification of Cys145-underivatized peptides as a function of time in HDX-MS measurements. Peptides 141-150 and 142-150, deriving from the cleavage of M^pro^ by pepsin during HDX-MS analysis, are quantified at different time points (0.25, 1, 5, 30, 120min) in the presence or absence of PPACK. The experiment was performed in triplicate. Each dot represents the results obtained for a biological replicate, with the box plot boundaries indicating the quantiles Q1 (25%) and Q3 (75%) and the median value denoted by a line across the box.

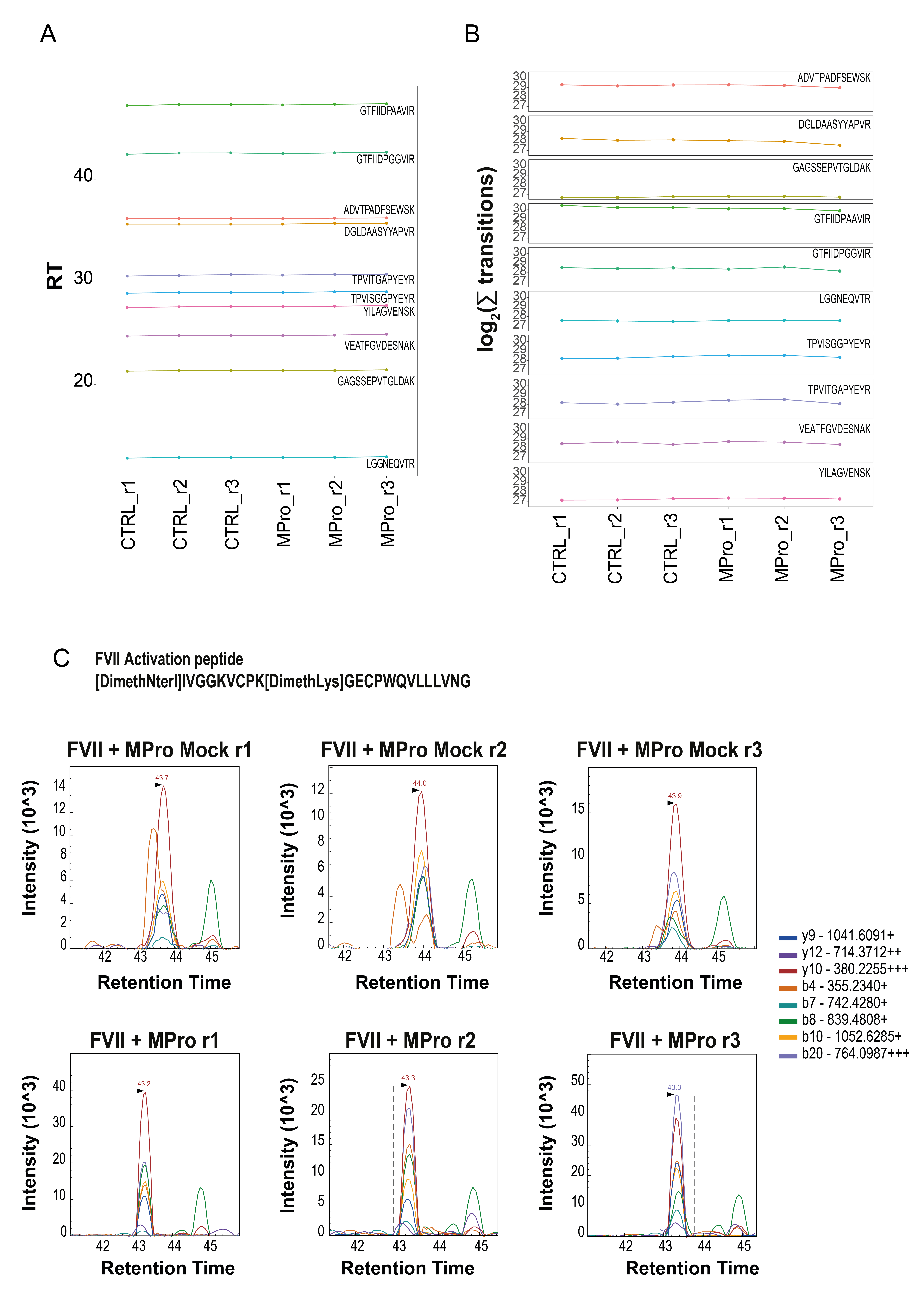

**Supplementary Figure S8. Targeted mass spectrometry analysis of FVII activation peptides. (A,B)** Analysis of iRT peptides used as an internal standard for the targeted analysis. Retention time and the integration of MS2 fragments are reported in panel A and B. (**C**) Elution profile of FVII activation peptide (Dimethyl)IVGGKVCPK(Dimethyl)GECPWQVLLLVNG. The top 8 most intense MS2 fragments are reported.
